# Tracing Neanderthal ancestry patterns through successive population expansions in Europe

**DOI:** 10.64898/2026.02.20.706963

**Authors:** Alexandros Tsoupas, Claudio S. Quilodrán, Jérémy Rio, Mathias Currat

## Abstract

During their expansion out of Africa, modern humans interbred with Neanderthals, leading to the introgression of Neanderthal DNA into their genomes. This initial dispersal created, in Europe, a Southeast–to–Northwest gradient of Neanderthal ancestry in European populations, which was preserved through later Neolithic migration. Here, we investigate the population dynamics that created and maintained this gradient across two successive expansions. We developed a three-population layered simulation framework to track the spatiotemporal evolution of Neanderthal ancestry under a model of interactions between Neanderthals, Palaeolithic hunter-gatherers, and Neolithic farmers. Our results indicate that the Neanderthal ancestry cline was shaped by the direction of the early hunter-gatherer expansion, the northern limit of the Neanderthal range, and strong reproductive isolation between lineages. We estimated that admixture between hunter-gatherers and Neolithic populations was an order of magnitude higher than that between Neanderthals and modern humans. These findings demonstrate how spatiotemporal analyses of ancient DNA provide insights into the dynamics and interactions of ancestral populations.

## Introduction

Modern humans (MH, *Homo sapiens*) originated in Africa between ∼300,000 and 100,000 years before present (YBP) (*1*). Between ∼60,000 and 40,000 YBP, MH populations expanded out of Africa into Eurasia, via the Middle-East (*2*). During this major dispersal, they encountered Neanderthals (NE), a related lineage that diverged from MH approximately 520,000–630,000 years ago (*3*). Neanderthals occupied a broad geographic range across Eurasia, from the Iberian Peninsula (*4*) to the Altai Mountains (*5*), until their extinction ∼40,000 YBP (*6*), following several millennia of coexistence with MH (*6*, *7*). The successful sequencing of the Neanderthal genome has showed that hybridization occurred between the two lineages during this period, with MH populations outside sub-Saharan Africa still carrying Neanderthal-derived DNA segments today (*8*).

On average, contemporary non-African MH populations harbor ∼2% Neanderthal ancestry in their genomes (*3*), with individual levels ranging between 1–4% (*9*). It is assumed that after an initial rapid decline due to purifying selection (*10*, *11*), only a limited number of genomic regions were subject to selection due to their association with specific traits or diseases (e.g. *3*, *12*–*14*). In contrast, the long-term stability of Neanderthal ancestry levels in Eurasian populations over the past 40,000 years suggests that most introgressed DNA persisted under neutral evolution (*9*), reflecting demographic rather than selective processes. Indeed, Neanderthal ancestry in Eurasian populations exhibits a spatial gradient that increases with distance from the Near East, consistent with admixture occurring during the initial expansion of MH out of Africa (*15*). This is supported by spatially explicit simulations, which have shown that when limited admixture occurs between a resident population at demographic equilibrium and an expanding population, a gradient of introgressed local alleles is expected to form along the expansion axis (*16*). Moreover, a recent study suggests that this cline is expected to halt near the edge of the hybridization zone, after which the levels of introgression stabilize (*17*). Such introgression gradients can arise solely from demographic and migratory processes, without invoking selection as a driving factor (*18*).

It has recently been suggested that the cline of Neanderthal ancestry was established during the initial settlement of Europe by early MH and has persisted to the present (*15*). However, the long-term persistence of such a cline is surprising given that successive episodes of migration, population partial turnover with admixture have profoundly reshaped the genomic composition of European populations throughout Prehistory (e.g. *19*–*21*). In particular, a major genomic change occurred with the emergence of agriculture in the Near East, which began spreading into Europe around 8,500 YBP and progressively replaced the hunter-gatherer lifestyle (*22*). This process, known as the Neolithic transition, was driven by the migration of early farming populations (FA) from Western Anatolia and the Aegean basin (*21*) into Europe (*23*, *24*). As these farming groups dispersed across the continent, they increasingly admixed with local hunter-gatherer (HG) populations (*25*), resulting in genomes that combined both FA and HG ancestries (*26*), with marked regional variation (*27*). It has been shown that the average level of Neanderthal ancestry in European populations declined during the Neolithic transition, driven by the arrival of early FA who carried slightly lower levels of Neanderthal ancestry than HG (*15*). Counterintuitively, the spatial gradient observed in HG persisted in the post-Neolithic populations, although the overall amount of Neanderthal ancestry declined. The demographic processes responsible for reducing the amount of ancestry while maintaining the gradient remain unexplored.

Here, we investigated the population dynamics factors underlying the creation and persistence of spatiotemporal patterns of Neanderthal ancestry in two ancient European populations: the pre-Neolithic HG and the subsequent Neolithic FA populations. To reproduce the gradient of Neanderthal ancestry observed in HG and its evolution following the Neolithic transition, we developed an original three-populations, spatially explicit simulation framework, based on the program SPLATCHE3 (*28*). This approach enabled us to explore how Neanderthal ancestry developed across Europe through space and time under the influence of two successive population expansions with admixture, each with distinct demographic, migratory, and interaction parameters. For each investigated scenario, we used an Approximate Bayesian Computation (ABC, *29*) to compare simulated outcomes with Neanderthal introgression estimates derived from a large published paleogenomic dataset of 715 European HG and FA individuals. Then, we jointly estimated the population dynamics and interaction parameters between Neanderthals and HG as well as between HG and FA. This enabled a direct comparison of population dynamics between these two key epochs of population turnover with admixture.

## Results and discussion

### Modern human arrival shaped the Neanderthal ancestry gradient

The large paleogenomic dataset, against which we compared the results of our simulations, is made up of 102 Paleolithic HG and 613 Neolithic FA (Fig. 1), retrieved from the AADR database (*30*). They shared on average 2.14% (standard deviation, sd = 0.38%) of their genome with Neanderthals. More specifically, Paleolithic HG have a mean proportion of Neanderthal ancestry estimated to 2.42% (sd = 0.42%) and Neolithic FA to 2.08% (sd = 0.34%), the latter being statistically significantly lower than the former (two sample *t*-test *p*-value = 6.2×10^−11^). Importantly, Neanderthal ancestry in Paleolithic HG displays a statistically significant gradient in populations, increasing from the Southeast to the Northwest of the continent (Table 1), confirming a previous observation (*15*). This gradient of Neanderthal introgression is expected under a model of MH range expansion with admixture (*31*). To assess whether the orientation of the observed gradient of Neanderthal ancestry in HG reflects the distribution of Neanderthals and the source of expansion of MH, we simulated four scenarios with a northern limit of the Neanderthal range varying between 55°N and 70°N (fig. S1). This range reflects the most plausible maximum latitude of Neanderthals in Europe (*32*, *33*), while their southern range extended until the Mediterranean and the Levant (*6*, *34*). We then performed model choice against the observed Neanderthal introgression gradient using an ABC framework (see Methods for details). All four investigated scenarios were able to reproduce to some extent the observed spatiotemporal gradient of Neanderthal ancestry in HG, as indicated by marginal density *p*-values > 0.05 (table S1). However, the observed gradient is most likely explained by the scenario lat55 for which the maximum range is located at latitude 55°N, which has a posterior probability of 56% (table S2). This is the only scenario that can be reliably identified, in 65.5% of the cases (fig. S2). In contrast, scenarios lat60 and lat65 are frequently misclassified as either lat55 or lat70, while lat70 is correctly identified in only 31.8% of the cases and is almost as often misclassified as lat55. Note that the confusion matrix indicates that when a simulation is classified as lat55, there is a 62% probability that it originates from a scenario where the Neanderthal range extended further north. Overall, the higher the latitudinal limit that was simulated, the less probable it was for the corresponding scenario to produce a spatiotemporal gradient with the same orientation as the observed one (table S3).

**Figure 1.**
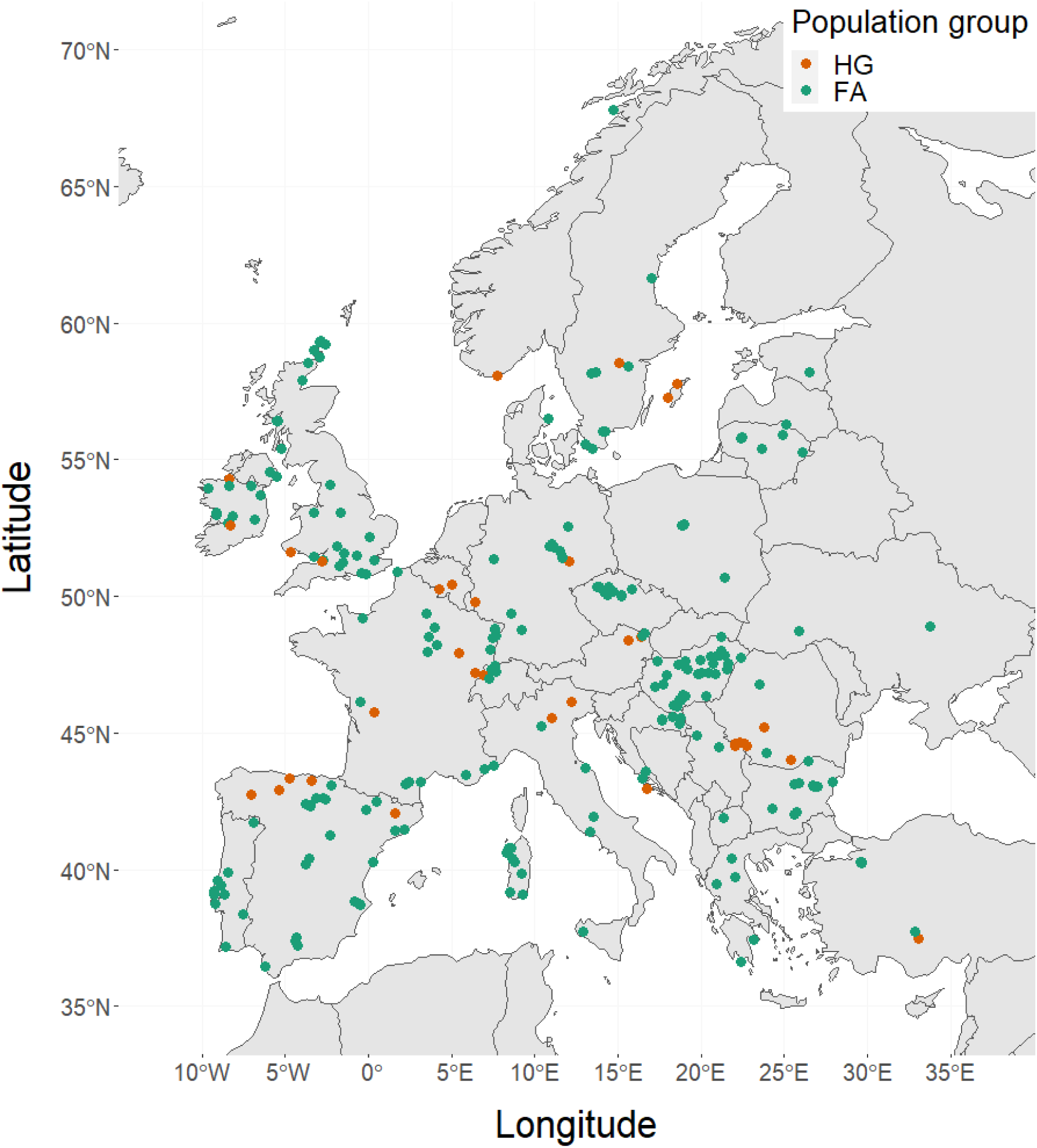
Geographic locations of the 715 individuals included in the analyses. Point colors indicate population group: red for hunter-gatherers (HG, *n*=102) and green for farmers (FA, *n*=613). Overlapping points occur where sampling locations are in close proximity.

**Table 1.**
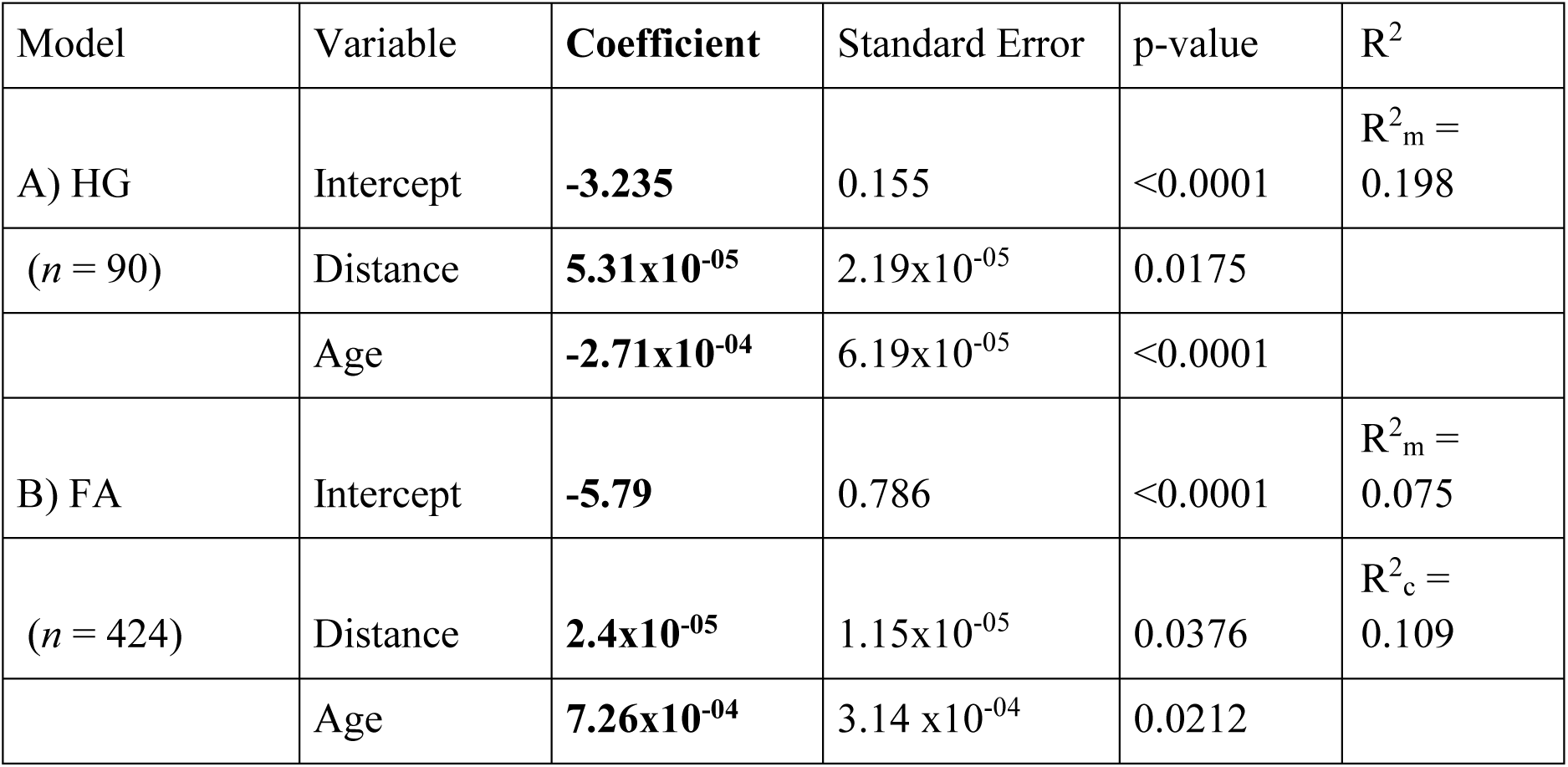
Statistical models describing Neanderthal ancestry levels in European Paleolithic hunter-gatherers (HG) and Neolithic farmers (FA). The coefficients of the linear models (in bold) were used as summary statistics for the ABC estimations. Marginal (R^2^_m_) and conditional (R^2^_c_) coefficients of determination indicate the variance explained by fixed effects alone and by both fixed and random effects, respectively. The response variable was log-transformed to ensure normally distributed residuals. Formula of the HG model called in the lm() function in R: *log(α_f4_) ∼ distance + age*. Formula of the FA model called in the lme() function of the nlme package in R: *log(α_f4_) ∼ distance + age, random=∼period | period, correlation=corLin(form=∼distance, nugget=F)*.

Our simulations are thus able to reproduce the observed spatiotemporal pattern of Neanderthal ancestry in HG most accurately when the northern limit of Neanderthal range was set at 55° latitude. Strikingly, this estimate, derived solely from paleogenomic data, corroborates archaeological and chronological evidence as well as habitat suitability models, which place the northernmost extent of Neanderthal presence at a similar latitude (*32* and references therein) or slightly farther south at ∼50° latitude (*35*). Although sporadic or low-density occupations may have occurred farther north, for example in parts of southern Scandinavia, which appear environmentally suitable for Neanderthal habitation (*32*), no definitive evidence of Neanderthal presence has yet been found north of 55°N in Europe (*36*). This northern limit likely would represent the maximal extent of stable Neanderthal occupation. Indeed, from ∼44 kya onward, Neanderthals progressively retreated from their northeastern range (*37*), with the contraction accelerating after ∼38 kya (*35*), ultimately leading to increasingly fragmented and isolated populations (*38*).

### The Neanderthal ancestry gradient was maintained during the Neolithic

As mentioned earlier, Neanderthal ancestry in Neolithic FA is statistically lower than in HG, but still distributed along a statistically significant Southeast-Northwest population gradient. Because previous simulation studies have shown that genetic gradients can occur not only parallel to an expansion axis but also perpendicular to it, through genetic surfing (*39*), we assessed whether the empirical Neanderthal ancestry clines observed in HG and FA could be obtained by multiple alternative demographic scenarios leaving similar signatures. We thus simulated four spatially explicit scenarios by alternatively varying the source of HG and FA expansions, placing them either in the Southeast (Northeast Africa for HG and Anatolia for FA) or the Southwest (Iberia for both HG and FA). In these descriptive simulations, the expansion origins of both HG and FA were arbitrarily defined, the goal being to approximate the general direction of their putative dispersal axis. First, we investigated whether the observed Southeast–Northwest gradient of Neanderthal ancestry reflects the directional expansion of early MH. Our simulations revealed that, regardless of the assumed source (Southeast or Southwest), HG populations expanding into Europe consistently developed a spatial gradient of Neanderthal ancestry increasing with distance from their origin (Fig. 2). This pattern appeared in 90–96% of simulations (Table 2). In the two scenarios where the HG expansion originates in the Southwest (S2-WW and S4-WE), a Southeast–Northwest gradient emerges in only 3% of simulations. It means that the observed Southeast–Northwest gradient is compatible only with scenarios where HG expanded from the Southeast (Scenarios S1-EE and S3-EW, Table 2), in line with archaeological evidence for MH dispersal into Europe from Southwest Asia (*40*, *41*). Thus, our descriptive simulations demonstrate that the Paleolithic expansion of HG into Europe is a strong candidate for establishing the observed Southwest–Northeast gradient of Neanderthal ancestry. Note that while the observed gradient indicates the broad direction of the expansion (*15*, *17*), it cannot pinpoint its exact source, which may lie in unstudied or poorly sampled regions.

**Figure 2.**
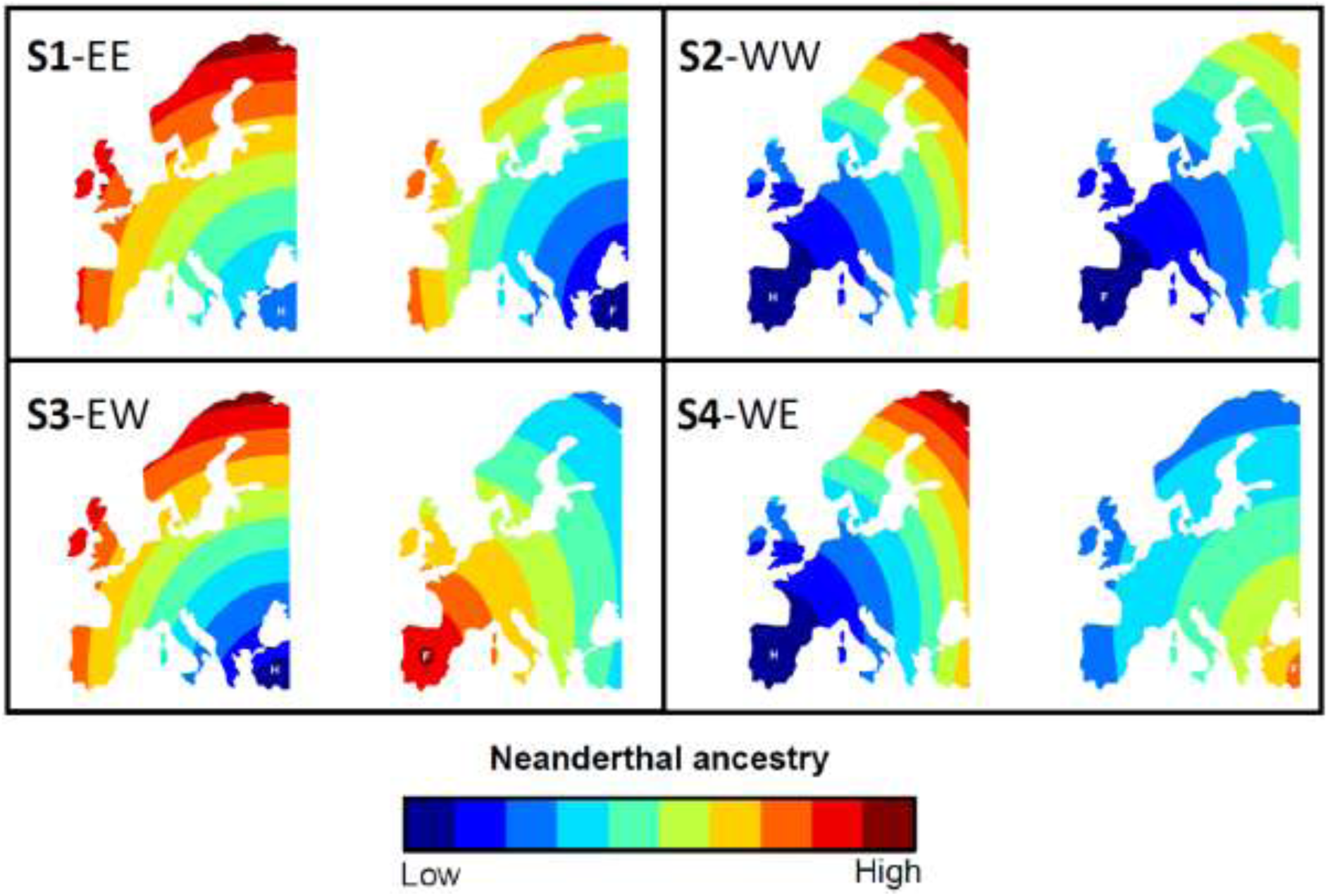
Extrapolated Neanderthal ancestry levels in HG and FA under four simulated expansion scenarios (S1–S4). Extrapolation was based on a linear model for HG and a linear mixed model for FA, fitted to the mean Neanderthal ancestry levels of each scenario. In each panel, the first map shows HG ancestry and the second map FA ancestry. White letters mark the reference point from which distances were measured (H = HG, F = FA). For FA, this always coincides with their expansion origin. For HG, it coincides with their own expansion origin in panels S2-WW and S4-WE, but with the FA origin in panels S1-EE and S3-EW, to allow distance to be calculated from a common reference point for both populations, as the source of HG in Northeast Africa is not represented in the map.

**Table 2.** Proportions of simulations producing a gradient of Neanderthal (NE) ancestry increasing with distance either from the respective expansion source or Anatolia (the presumed entry point of HG and FA into Europe). Results are shown separately for HG and FA under four expansion scenarios: S1(EE) both groups expand from their presumed origins (HG from Africa, FA from Anatolia); S2(WW) both expand from a hypothetical location in central Iberia; S3(EW) HG expand from Northeast Africa and FA from central Iberia; S4(WE) HG expand from central Iberia and FA from Anatolia. For each scenario, we also report the proportion of simulations in which HG exhibit higher mean Neanderthal ancestry than FA.

| Scenario | Percentage of simulations |  |  |  |
| --- | --- | --- | --- | --- |
|  | S1-EE | S2-WW | S3-EW | S4-WE |
| NE ancestry cline increasing in HG with distance from the HG source | 0.90 | 0.96 | 0.90 | 0.96 |
| NE ancestry cline increasing in FA with distance from the FA source | 0.92 | 0.95 | 0.11 | 0.06 |
| NE ancestry cline increasing in HG with distance from Anatolia | 0.90 | 0.03 | 0.90 | 0.03 |
| NE ancestry cline increasing in FA with distance from Anatolia | 0.92 | 0.05 | 0.88 | 0.06 |
| Higher NE ancestry in HG than in FA | 0.57 | 0.96 | 0.07 | 0.37 |

Furthermore, our spatially explicit simulations show that the pattern of Neanderthal ancestry in FA populations, acquired through admixture with HG, depends not only on the origin and direction of the FA expansion, but also on those of HG. While the HG expansion defines the orientation of the gradient, the FA expansion determines whether it increases or decreases. When FA expanded along the same axis as HG (Fig. 2A–B), Neanderthal ancestry in FA increased with distance from their expansion source, a pattern reproduced in 92–95% of simulations (Table 2). By contrast, when FA expanded orthogonally to the HG expansion (Fig. 2C–D), the gradient reversed, with ancestry decreasing with distance from the FA source (89–94% of simulations).

Since in this case the FA source is in the Iberian Peninsula, the ancestry decreasing with distance from it means that the ancestry is increasing with distance from Southeast Europe. Consequently, the observed increase of Neanderthal ancestry in FA with distance from the Southeast is only compatible with the two scenarios in which HG themselves expanded from the Southeast (S1-EE and S3-EW, Table 2), regardless of where the FA expansion began. This finding aligns with François *et al*. (*39*), who showed that successive expansions can maintain the spatial genetic pattern established during the initial colonization, provided that gene flow between populations is sufficient. In the case of Neanderthal ancestry, the similarity between pre- and post-Neolithic patterns reflects the FA populations “inheriting” an already established gradient. This highlights the importance of attributing observed gradients to the demographic events that originally generated them, rather than to subsequent population movements (*42*). Overall, our simulations indicate that the Neanderthal ancestry gradient was created by the initial HG expansion and subsequently maintained during the subsequent FA dispersal with admixture.

### The temporal fate of Neanderthal ancestry in European populations

A central question regarding Neanderthal introgression in MH is its temporal fate. The initial proportion of Neanderthal DNA in MH could have been as high as ∼6–10% and declined rapidly within the first 10–20 generations after admixture, due to purifying selection against weakly deleterious Neanderthal alleles (*10*, *11*). These alleles would have been efficiently purged from the MH gene pool because of their larger effective population size, leading to a rapid stabilization around present-day levels. The paleogenomics dataset we analyzed nevertheless revealed a small but statistically significant decline of Neanderthal ancestry in HG (Table 1). During the investigation of the northern limit of Neanderthal habitat, we simulated a scenario excluding the Neolithic expansion (lat55) and, interestingly, these simulations of neutral loci replicated this slight temporal decrease, without considering any selective pressure (table S3). This suggests that genetic drift within structured HG populations can account for a slow gradual reduction of Neanderthal ancestry without invoking long-term selection after the initial purging phase.

However, our simulations suggest that the markedly lower Neanderthal ancestry observed in post-Neolithic Europeans compared to pre-Neolithic groups is more consistent with the arrival of FA and their admixture with HG, than with drift alone. This is supported by the fact that the scenario with the Neolithic expansion had a posterior probability of 79%, when using ABC to compare it to a scenario without the Neolithic expansion.

All four simulated scenarios revealed an expected statistically significant difference in Neanderthal ancestry between HG and FA (Welch’s *t*-tests, table S4). Two scenarios show more Neanderthal ancestry in FA than in HG, while two show the reverse. In S1-EE the mean difference is 0.007 in favor of HG; in S2-WW, 0.026 in favor of HG; in S3-EW, 0.021 in favor of FA; and in S4-WE, 0.009 in favor of FA. Our descriptive simulations thus show that the starting location of the FA expansion is critical to explain the decrease of Neanderthal ancestry in post-Neolithic populations because this expansion needs to start with a population having a lower-than-average amount of Neanderthal introgression. If the FA expansion had begun in a region where Neanderthal ancestry was higher compared to other regions, then the mean difference in Neanderthal ancestry between HG and FA would have shifted in favor of the FA (e.g. scenarios S3-EW and S4-WE). More importantly, among the two scenarios that reproduce the observed Southeast–Northwest cline (S1-EE and S3-EW), only S1-EE, where HG and FA expand in the same direction, resulted in higher Neanderthal ancestry in HG than in FA, reproducing the observed pattern. This indicates that the Neolithic expansion drove the overall decline in Neanderthal ancestry levels while maintaining its gradient with the same orientation. This result is consistent with the origin of migrating early FA in the Aegean area and western Anatolia (*21*), where pre-Neolithic populations carried less Neanderthal ancestry than their contemporaries in Central Europe (*43*, *44*). Interestingly, Neolithic FA display an opposite temporal trend than HG, with a slow statistically significant increase in Neanderthal ancestry (Table 1), as also observed in Quilodrán *et al*. (*15*). This increase can be explained by the increasing admixture between HG and FA during the Neolithic transition (*25*). Because HG carried higher proportions of Neanderthal DNA, their gradual admixture with FA led to a slight increase of Neanderthal ancestry in FA.

Overall, although our scenarios are extreme simplifications of the much more complicated genomic and migratory history of ancient humans (*27*), our simulations demonstrate that two consecutive population expansions along a Southeast–Northwest axis have the potential to reproduce both the observed clinal pattern of Neanderthal ancestry in European populations since the Paleolithic, and the continent-wide reduction of Neanderthal ancestry during the Neolithic. Under the proposed model, this decline occurred through dilution, as initially proposed by Vernot and Akey (*45*), and may account for the lower levels of Neanderthal ancestry observed in present-day Europe compared to East Asia (*46–48*), assuming a similar process did not take place in Eastern Eurasia.

### Population dynamics parameter estimation

Based on the scenario which produces more often the observed NE ancestry populational patterns (gradients and difference between HG and FA, Table 2), we performed a parameter estimation using ABC. This scenario represents two successive expansion sources in the Southeast (scenario S1-EE) and the most probable northern limit of the Neanderthal range (55° latitude, table S2). The ABC analysis indicated that our three-population layered model was able to jointly reproduce the observed spatiotemporal Neanderthal ancestry patterns in FA and HG. It yielded a marginal posterior density *p*-value of 0.69 for a tolerance rate δ = 0.01 (*p*-value = 0.71 for δ = 0.005 and *p*-value = 0.68 for δ = 0.03), thus allowing us to properly estimate key population dynamics parameters.

Fig. 3 shows a graphical representation of the prior and posterior distributions of each parameter with δ = 0.01 and table S5 shows the associated posterior distribution characteristics. The hybridization rate between Neanderthals and HG (*γ_H_*) is characterized by a well-defined (Fig. 3A) and narrow posterior distribution (95% HDI 0.0038-0.008), with a mode of ∼0.006 and a low relative bias (0.088). This shows that the observed gradient of Neanderthal ancestry in HG developed under strong reproductive isolation between Neanderthals and early MH, with less than 0.9% of their contacts producing viable, fertile offspring, in accordance with estimates obtained from modern genomic data using a similar spatially explicit simulation approach (0.5%-0.8%, CI between 0.33% and 1.3%) (*49*, *50*). The ratio of effective population size of HG over Neanderthals (*Kratio_HN_*) is estimated to 7.7 (Fig. 3C) and characterized by a large relative bias (0.48) and broad posterior distribution (95% HDI between 2.3 and 9.7, table S5). This estimate is consistent with a much smaller effective population size generally inferred for Neanderthals compared to MH (*3*, *51*). Neanderthal effective sizes are typically estimated around ∼3,000-3,500 individuals (*3*, *52*, *53*), and may have been up to 20% lower when accounting for reverse introgression from MH, that possibly took place ∼250,000-200,000 and ∼120,000-100,000 years ago (*54*). In contrast, MH effective population size is generally estimated at 10,000–15,000 individuals (*55*).

**Figure 3.**
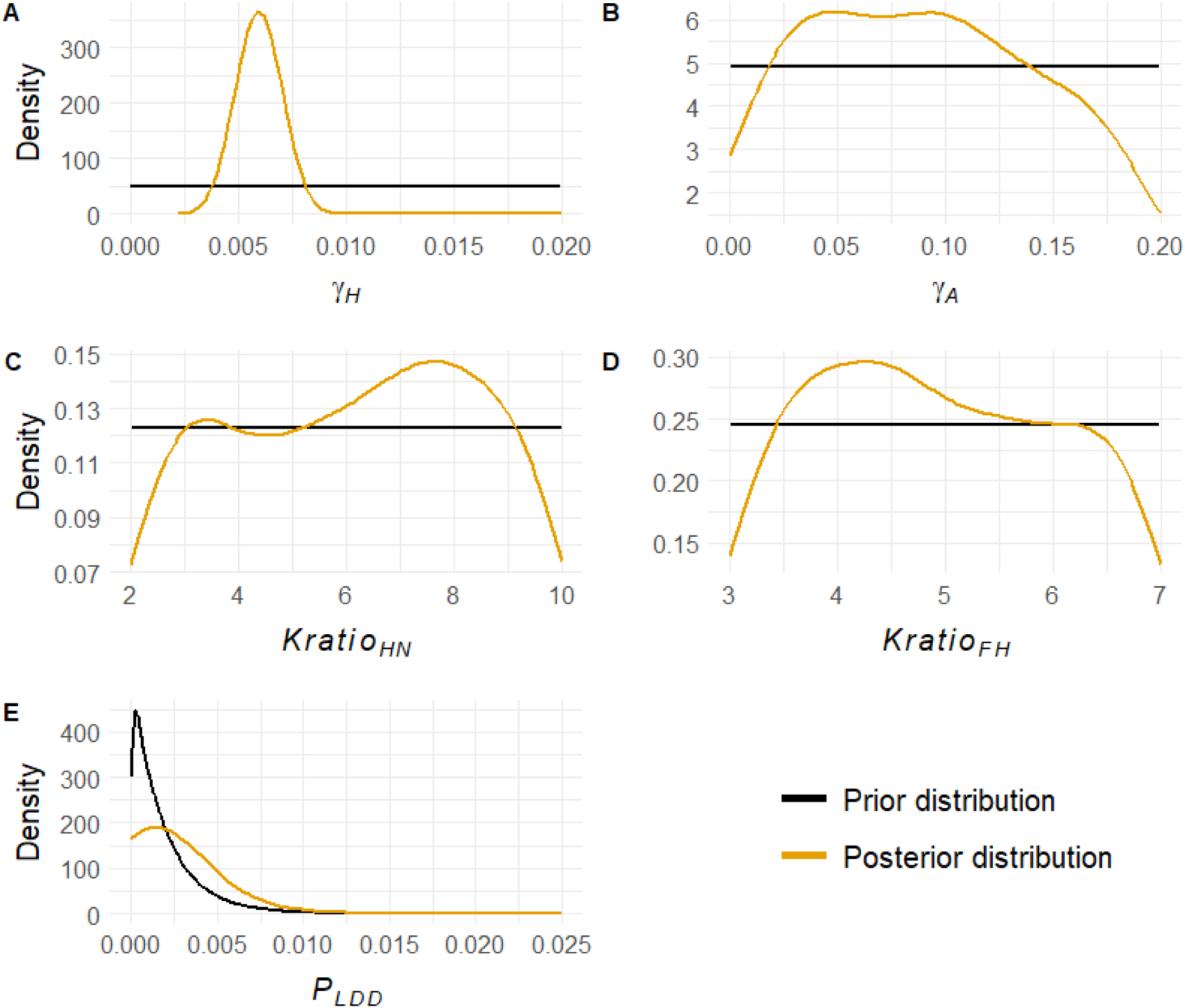
Prior and posterior distributions of the estimated parameters. The black line shows the prior distribution used in the simulations, and the yellow line the posterior distribution inferred from the ABC estimation. (A) Hybridization rate between Neanderthals and HG (*γ_H_*); (B) Assimilation rate of HG into FA (*γ_A_*); (C) Ratio of HG to Neanderthal effective population size (*Kratio_HN_*); (D) Ratio of FA to HG effective population size (*Kratio_FH_*); (E) Proportion of FA migrations occurring as Long Distance Dispersals (*P_LDD_*).

The ABC parameter estimation identified key factors required to maintain the observed Neanderthal ancestry gradient in FA. The estimated assimilation rate of HG into FA (*γ_A_*) is 0.049, but quite imprecise, with a broad 95% HDI (0.002–0.181; table S5), and a high relative bias (1.93). Nevertheless, the point estimate is very close to the ∼0.046 reported by Tsoupas *et al*. (*25*) based on molecular diversity computed on 67 high quality paleogenomes. It gives an order of ∼5% of contacts between HG and FA resulting in gene flow between them. This limited gene flow may reflect ecological, economic, or cultural segregation (*56*). Notably, the HG–FA assimilation rate is estimated to be ∼15 times higher than the Neanderthal–HG hybridization rate (table S5), in accordance with the strong interspecific reproductive barriers between Neanderthals and MH, which obviously do not exist between HG and FA.

Furthermore, we estimated the effective population size of FA to be, on average, four times larger than that of HG (*Kratio_FH_*= 4.25, Fig. 3D), with a relatively low bias (0.22) and quite large posterior distribution (95% HDI = 3.14-6.82, table S5). This is very close to the estimates (∼5) by Tsoupas *et al*. (*25*), which was based on paleogenomic pseudo-haploid nucleotide diversity rather than genomic ancestry proportions. This ratio in favor of FA is in accordance with the lower genetic diversity measured in HG compared to FA populations (*57*). It is noteworthy that under our modelling framework, the estimated population size ratio between HG and FA is almost half of that between HG and NE, with 68% probability of the latter being larger than the former, which reflects the particularly small effective size reported for late Neanderthals (*38*).

The proportion of long-distance dispersals (LDD) during the Neolithic expansion could not be precisely determined but falls within very low values (<0.8%), with zero included in the credible interval. The posterior distribution has a mode of 0.0015, a high relative bias (1.81), and a posterior 95% HDI of 0–0.0078 (table S5). Additionally, a set of supplementary simulations, using the same simulation framework and parameter priors but without simulating LDD, was able to replicate the observed Neanderthal ancestry gradient in FA, as the marginal density *p*-value of the simulations was 0.68, which means that the observed gradient can be formed even in the absence of LDD. These results indicate that LDD had minimal impact on the gradient and overall levels of Neanderthal ancestry in farming populations, even though it has been shown to reduce introgression (*58*). LDD may have played a more substantial role in increasing the speed of the FA expansion, as suggested by Tsoupas *et al*. (*25*).

### Model strengths, limitations and perspectives

The strength of our approach lies in its spatially explicit, three-layered design applied to an extensive paleogenomic dataset. This framework enabled us to define spatiotemporal patterns of Neanderthal ancestry and use them for demographic inference. By incorporating introgression from a third population, our framework extends the previous version of SPLATCHE3 (*28*), which was restricted to two contemporaneous lineages. This allowed us to model Neanderthal ancestry across space and time through two successive expanding populations with admixture, Paleolithic HG and Neolithic FA, and to compare alternative scenarios against a large paleogenomic dataset in order to identify the processes underlying the observed cline of Neanderthal ancestry in Europe.

While powerful, our framework is computationally demanding and rests on simplifying assumptions, some of which warrant further investigation. For instance, we assumed constant demographic and migratory parameters for Neanderthals, HG, and FA. Future work should examine the impact of regional heterogeneity (*59*, *60*) and fluctuating population sizes (*61–63*). Although our simulations suggest minimal Neanderthal presence north of 55° latitude, this pattern could potentially be refined by modelling variable population densities, e.g. larger in the south, lower in the north, to investigate whether it fits better the observed ancestry patterns. Because our framework currently considers neutral loci, it likely underestimates the initial level of admixture, before the purging of Neanderthal DNA during the first 10–20 generations (*10*, *11*). Incorporating this effect would need to include selection parameters in the model, which would add another level of complexity. Likewise, the model cannot address whether reproductive isolation between Neanderthals and MH was pre- or post-zygotic. For the sake of simplicity, we also assumed a constant HG–FA assimilation rate, despite evidence that it increased during the Neolithic transition (*25*, *64*), and did not account for asynchronous or multi-wave expansions. Also, our model does not include minor or dead-end MH migrations that left little genetic trace in later populations (*65*).

Lastly, an important event not included in our model, but warranting further investigation, is the Last Glacial Maximum (LGM, 19–27 kya), a period during which the fate of human populations remains poorly understood. For example, it is unclear to what extent HG retreated to southern refugia and whether their distribution was continuous (e.g. *59*) or fragmented (e.g. *66*, *67*). This period is particularly relevant because it falls between the two events examined here: the initial settlement of Europe by HG and the Neolithic transition. Understanding how Neanderthal ancestry gradients persisted or were reshaped after the LGM would provide critical insights.

## Conclusion

Our original simulation framework represents a step forward toward developing more detailed scenarios for investigating how demographic variation, migration, and population interactions have shaped spatiotemporal patterns of genomic ancestry. As paleogenomic data continues to accumulate with improved sequencing quality, this iterative approach has the potential to substantially refine our understanding of ancient population dynamics and enable more precise demographic inferences about major events that marked the history of our ancestors across different periods. Our study highlights the key role of demographic processes during range expansions in shaping patterns of archaic introgression, providing alternative explanations to selection or uneven hybridization, though these mechanisms are not mutually exclusive. More broadly, our study demonstrates the potential of spatiotemporal ancestry patterns to reconstruct expansion directions and to estimate demographic parameters such as admixture rates and relative population sizes. Strikingly, our results show that interspecific introgression can also illuminate intraspecific demographic processes. The framework can be readily extended to later periods of Prehistory, such as the Bronze Age steppe migrations (*19*, *20*), or modified to study introgression from other archaic groups like Denisovans, as well as to trace the genomic composition of human populations through time. Beyond the case studied here, this approach offers broad applicability for investigating species interactions across diverse research contexts.

## Material and Methods

### Three population layered spatially explicit simulations

We used SPLATCHE3 (*28*) to simulate interactions between Neanderthals and MH and the resulting patterns of Neanderthal ancestry. The program models two overlapping population layers on a spatially explicit ASCII map divided into cells, each containing two demes with distinct demographic parameters. Layers interact through user-defined settings. Simulations proceed forward in discrete generations for demography, followed by a backward coalescent step that generates molecular diversity in samples from various locations and times, as defined by the user.

More specifically, we used a modified version of SPLATCHE3 (published in *25* and available on Zenodo) that includes three new features. First, we extended the original two-population framework to three populations (A, B, C), with the condition that only two coexist at any given time. Once one of the initial populations (A) goes extinct through competition with the other (B), a third population (C) can be introduced into the empty layer and interact with the surviving one (B). In this study, these correspond to Neanderthals, HG, and FA: Neanderthals and HG were the initial populations, while FA appeared later, derived from HG in Anatolia after Neanderthal extinction. SPLATCHE3 then outputs the proportion of loci in HG and FA samples that originated from Neanderthals (i.e. ancestry proportion). These proportions represent the same measure as the F4-ratio (*68*), which was the statistic used to estimate Neanderthal genomic contribution to MH in the observed dataset. Second, we allowed admixture models to change at a predefined generation, switching from hybridization (Neanderthal–HG) to assimilation (HG–FA). In the hybridization model, contacts generate hybrid individuals assigned to one parental species, whereas in the assimilation model, gene flow goes from HG into the FA population. Gene flow frequency is controlled by a parameter *γ*: in the hybridization model, *γ_H_* represents the proportion of individuals producing hybrid offspring in the next generation; in the assimilation model, *γ_A_* represents the proportion of individuals from one population incorporated into the other (see SPLATCHE3 manual, www.splatche.com). Third, competition between layers could switch from density-dependent (*16*, *42*) to a user-specified rate (*α*). The intensity of competition determines the duration of cohabitation. Because the duration of Neanderthal–HG coexistence in Europe is uncertain (*6*, *7*), we modelled their competition as density-dependent, which simplified simulations without affecting HG sampling. For HG–FA interactions, however, density-dependent competition caused HG to disappear too early, preventing dataset recovery. Cohabitation time being primarily determined by population sizes and *α*, we set *α* to a fixed value during the early FA expansion to allow prolonged coexistence, then switched back to density-dependent competition 10 generations after the last HG sample, ensuring HG extinction during the Neolithic. To maximize simulation success, *α* was estimated individually for each simulation relative to the ratio of FA to HG carrying capacity (*Kratio_FH_*). The procedure was as follows: (1) 200,000 initial simulations were run, and only those reproducing the observed dataset (correct samples in time and space) were retained; This corresponds to the “demographic filter” in Rio *et al*. (*68*); (2) *Kratio_FH_* and *α* from the retained simulations were used to fit a linear model (*α* ∼ *Kratio_FH_*) with the lm() function in R; (3) *α* for each subsequent simulation was predicted from its *Kratio_FH_* using predict.lm(). This yielded *α* values between 0.113 and 0.267, very close to the one estimated in Tsoupas *et al*. (*25*) which used a paleogenomic dataset specifically designed to study the Neolithic transition along the Danube route (*α* = 0.15-0.327).

The simulation framework followed Quilodrán *et al*. (*16*), using a map of Europe divided into 75 × 53 cells of 100 × 100 km (fig. S3). The initial expansion sources were set at: (1) an arbitrary point in Northeast Africa (30°, 34°) for the HG layer, representing the out-of-Africa expansion following Quilodrán *et al*. (*15*); and (2) an arbitrary point in Anatolia (37°, 36°) for the FA layer, consistent with paleogenomic evidence for the European Neolithic expansion starting from this area (*21*, *66*). Sample coordinates were converted to WGS84 Plate Carree format using the proj4 R package (*69*). The lower-left map corner was set at 3201000, –1647000 (WGS84 Plate Carree). Simulations started at 70,000 YBP to allow Neanderthals sufficient time to populate Europe and reach demographic equilibrium before MH appeared. With a generation time of 25 years (*70*), this corresponded to 2,800 simulated generations. Because our focus was on MH expansions, earlier generations were omitted to improve computational efficiency. The HG expansion into Europe was set at generation 800 (∼50,000 YBP; *71*), and the FA expansion at generation 2,380 (∼10,500 YBP; *22*).

The mean ages of observed genomes were converted to generations since the start of the simulation. Constant parameters followed Quilodrán *et al*. (*16*), while Table 3 lists those that varied across simulations and their prior ranges. For each genome, we simulated 50 independent, neutrally evolving genomic loci to estimate Neanderthal ancestry. This number balances computational efficiency with accuracy (*68*).

**Table 3.** Prior value ranges of parameters that were varying between simulations and values of the constant parameter.

| Parameter | Prior distribution |
| --- | --- |
| Hybridization rate between NE and HG ($\gamma_H$) | 0.0-0.02 |
| Assimilation rate of HG into FA ($\gamma_A$) | 0.0-0.2 |
| NE effective population size ($K_{NE}$) | 200 |
| HG effective population size ($K_{HG}$) | 400-2,000 |
| FA effective population size ($K_{FA}$) | 1,200-14,000 |
| HG to NE effective population size ratio ($K_{ratio_{HN}}$), defined as $K_{HG}/K_{NE}$ | 2-10 |
| FA to HG effective population size ratio ($K_{ratio_{FH}}$), defined as $K_{FA}/K_{HG}$ | 3-7 |
| Proportion of Long Distance Dispersals ($P_{LDD}$) | 0-0.025 |

Interbreeding between Neanderthals and HG occurred in any cell of co-occurrence, with a probability defined by the hybridization rate (*γ_H_*). Assimilation of HG into FA demes was governed by the assimilation rate (*γ_A_*), which applied where both populations coexisted. These rates reflect the probability that contact between two populations results in gene flow. For *γ_H_*, the prior range was 0–0.02 and for *γ_A_*, 0–0.2 (*49*, *64*). Assimilation was unidirectional (HG into FA), and the switch from hybridization to assimilation was set one generation before the emergence of FA (generation 2,379).

The ratio of HG to Neanderthal effective population size (*Kratio_HN_*) ranged between 2 and 10 (*72*), while the ratio of FA to HG size (*Kratio_FH_*) ranged between 3 and 7 (*25*). For each simulation, both ratios were drawn from uniform distributions. Neanderthal effective size was fixed at 200 individuals, from which HG and FA sizes were derived: HG size = 200 × *Kratio_HN_*; FA size = HG size × *Kratio_FH_*. As SPLATCHE3 requires haploid counts, all effective sizes were doubled to obtain the number of gene copies used in the simulations.

Neanderthals and HG migrated in a 2D stepping-stone manner (*73*). FA also migrated primarily in a stepping-stone fashion, but with a small probability (*P_LDD_*) of long-distance dispersal events, ranging from 0 to 0.025. *P_LDD_* values were drawn from an exponential distribution to explore more often lower values, as Tsoupas *et al.* (*25*) showed that only a small percentage of the total migrations took place over long distances. Migration distances were sampled from a gamma distribution with shape *β* = 1.209 and rate *λ* = 0.15046, following Rio *et al*. (*68*).

### Paleogenomic dataset

Paleogenomic data were retrieved from the AADR database (Allen Ancient DNA Resource v50.0; *30*), which contained 10,391 genomes. We retained only European genomes (longitude < 34°) belonging to Paleolithic/Mesolithic hunter-gatherers (HG) or Neolithic/Chalcolithic farmers (FA). Group assignment was based on the AADR “GroupID” column, the mean age of the genome (genomes older > 15,000 years were all categorized as HG), or the original publication. For genomes represented by multiple versions, we kept the one with the highest SNP coverage. We filtered out putatively contaminated genomes by using the contamination warning estimated through Linkage Disequilibrium (*74*): Entries flagged as “Model_Misspecified” or “Very_High_Contamination,” or with “contam/possible.contam” in their GroupID, were excluded. We also excluded genomes lacking geographic coordinates. For dating, we used the mean age (YBP) provided in the database, corresponding either to the midpoint of the 95.4% CI calibrated radiocarbon date or the midpoint of the archaeological context range.

### Estimation of Neanderthal ancestry

Neanderthal ancestry was estimated using F4-ratios (*α_f4_*) as described by Reich *et al*. (75). The F4-ratio, defined as *α_f4_* = F4(A, O, X, C) / F4(A, O, B, C), measures the proportion of ancestry in a test genome (X) derived from population B, relative to a sister population C. Following Petr *et al*. (43), we used Dinka from eastern Africa as the unadmixed sister population (C), the Altai

Neanderthal as sister population (A) of Vindija Neanderthal (B), and chimpanzee as outgroup (O). These groups are available in the AADR database as “Altai_Neanderthal.DG,” “Vindija_Neanderthal.DG,” “Dinka.DG,” and “Chimp.REF.” F4-ratios were computed with ADMIXTOOLS 2 (76) and genomes with non-significant values (|Z| < 3) were excluded. We refer to the *α_f4_* as Neanderthal ancestry. After filtering, 715 genomes were retained (102 HG and 613 FA), ranging in age from ∼35,000 to ∼4,000 YBP. Genome metadata are provided in data S1, and sampling locations are shown in Fig. 1.

### Summary statistics used for the ABC analysis

The observed dataset was aggregated to match the simulated samples. Genomes from the same population group (HG or FA), sampled in the same simulated deme and generation (25-year intervals), were grouped and their Neanderthal ancestry averaged, yielding 514 population samples (90 HG, 424 FA; data S2). We then estimated the mean Neanderthal ancestry in the whole dataset and separately in HG and in FA, with these values being used as summary statistics in the Approximate Bayesian Computation (ABC; *29*) analysis (see below).

For each sample, geographic distance from the source of the corresponding expansion (fig. S3) was calculated with the gmt R package (https://rdrr.io/cran/gmt/). In FA, distance from the expansion source (*d_FA_*) was correlated with sample age in generations (Pearson’s r = 0.57, *p*-value < 2×10^−16^), with more distant samples tending to be younger. To account for this, FA samples were divided into three periods of ∼84 generations each, and Neanderthal ancestry was modeled as a function of *d_FA_* and age using a linear mixed model (LMM; Table 1). The period was included as a random slope and intercept, with linear spatial autocorrelation. LMMs were fitted with the *nlme* R package (*77*). In HG, distance from the expansion source (*d_HG_*) was not correlated with age (Pearson’s r = 0.06, *p*-value = 0.6), so Neanderthal ancestry was modelled with a simple linear model including *d_HG_* and age (Table 1). In both models, Neanderthal ancestry was log-transformed to ensure normally distributed residuals. The coefficients from these models (Table 1), along with mean Neanderthal ancestry in HG and FA, were then used as statistics in the ABC analysis.

### Impact of the northern limit of Neanderthal habitat on the Neanderthal ancestry gradient

We tested whether the northernmost limit of Neanderthal habitat in Europe influenced the observed Neanderthal ancestry gradient in HG. This analysis relied on ABC model choice to compare scenarios and assess their ability to reproduce the observed gradient. We simulated four scenarios with different northern habitat limits (fig. S1): 55° (“lat55”), 60° (“lat60”), 65° (“lat65”), and 70° (“lat70”). The 55° latitude reflects the generally accepted northern limit of NE occupation (*32*, *36*). Because adaptability to colder climates was not a limiting factor for their survival further north (*33*, *78*) and preservation issues cannot be ruled out (*79*), we explored the region north of 55° by dividing it into equal latitudinal bands of 5° each. Only Neanderthals and HG were simulated, with 400,000 runs per scenario. For each simulation, Neanderthal ancestry proportions in HG were fitted with the linear model described in Table 1A, and the resulting intercept, distance, and age coefficients, together with mean Neanderthal ancestry in HG, were used as summary statistics.

Goodness of fit between simulated and observed data was assessed with ABCtoolbox2 (*80*) by computing marginal density *p*-values under the null hypothesis that simulations matched the observed statistics. Euclidean distances were calculated, and a tolerance δ of 0.005, 0.01, or 0.03 (corresponding to 2,000, 4,000, or 12,000 retained simulations) was applied to test robustness to tolerance level.

Additionally, for each scenario we estimated the proportion of simulations which produced a gradient of NE ancestry increasing in HG with distance from the expansion source, the proportion of simulations which produced a gradient of NE ancestry in HG decreasing with time, and the proportion of simulations that produced both such gradients. This was done to assess whether the simulations could reproduce the overall observed trend and to determine which scenario most frequently generated a gradient similar to the observed one.

Model choice was performed with the abcrf R package (*81*), which implements a random forest approach with strong discriminative power and relatively low simulation requirements. The abcrf R package estimates the posterior probability of the scenario chosen by the most trees. We tested different forest sizes and retained 2,000 trees, as larger forests did not further reduce misclassification error.

### Impact of HG and FA expansion directions on the Neanderthal ancestry gradient

Because genetic clines may differ from the axis of population expansion (*39*, *82*), we examined the effect of the direction of the two consecutive MH expansions on the observed gradient of Neanderthal ancestry in Europe: the arrival of HG and the later migration of FA. Using descriptive simulations, we compared the gradients produced when both groups expanded from the Southeast (the presumed entry point of both MH and later FA into Europe) toward the Northwest, versus the gradients when one or both groups expanded from a hypothetical entry point in the Iberian Peninsula toward the Northeast. We performed 10,000 simulations for each of four scenarios differing in the expansion origins of HG and FA:

● **S1-EE:** Both HG and FA expand from the Southeast, HG from a chosen location in Northeast Africa and FA in Anatolia (fig. S4A).
● **S2-WW:** Both expand from the Southwest, in the Iberian Peninsula (fig. S4B).
● **S3-EW:** HG expand from the Southeast (Northeast Africa) and FA from the Southwest (Iberian Peninsula, fig. S4C).
● **S4-WE:** HG expand from the Southwest (Iberian Peninsula) and FA from the Southeast (Anatolia, fig. S4D).

For each scenario, we used simulated Neanderthal ancestry proportions to fit the same linear (HG) and linear mixed models (FA) applied to the observed data (Table 1). We then estimated, separately for HG and FA, the proportion of simulations of each scenario in which Neanderthal ancestry increased with distance from the respective starting location of each expansion or with distance from Anatolia (the presumed entry point of HG and FA into Europe) (Table 2). Finally, we estimated for each scenario the proportion of simulations in which HG had higher NE ancestry than FA (Table 2) and the mean simulated difference in Neanderthal ancestry between HG and FA (table S4).

### Population dynamics parameters shaping the Neanderthal ancestry gradient

We assessed whether levels of gene flow between Neanderthals and HG, between HG and FA, and the relative effective sizes of the three groups contributed to the observed Neanderthal ancestry gradient, as these factors are known to influence introgression patterns (*16*). We inferred the parameter values most compatible with the observed levels and gradient of Neanderthal ancestry by using 200,000 simulations of HG and FA expansions from arbitrarily chosen locations representing their presumed origins (Northeast Africa for HG, Anatolia for FA; fig. S3) with the three-population version of SPLATCHE3.

Parameter estimation was carried out with ABCtoolbox2, which computes the minimized Euclidean distance between simulated and observed summary statistics. The closest simulations are retained, and an ABC-GLM regression adjustment (*83*) is applied to their parameter values to estimate posterior distributions. As summary statistics, we used mean Neanderthal ancestry in HG and FA and the coefficients of the respective linear models (Table 1, bold), for a total of eight statistics.

The model fit was assessed as in the analysis of the latitude of Neanderthal habitat. To test robustness to the tolerance rate (δ), we retained 1,000, 2,000, and 6,000 simulations (δ = 0.005, 0.01, and 0.03, respectively). The posterior distribution of each parameter was compared with the respective prior, and posterior modes, means, and 95% Highest Density Intervals (HDIs) were estimated. In the main text, the posterior modes are used as point estimates. For the two population ratios (*Kratio_HN_* and *Kratio_FH_*), we also estimated a two-dimensional posterior distribution and used Markov chain Monte Carlo (MCMC) sampling to estimate the probability that *Kratio_HN_* is larger than *Kratio_FH_*.

To evaluate precision, we computed the relative bias of the posterior mean and mode. For each retained simulation, the simulated summary statistics were treated in turn as pseudo-observed values with known parameters, and posterior estimates were obtained from the remaining simulations through an ABC estimation. Relative bias was then calculated as: 𝑟𝑒𝑙𝑎𝑡𝑖𝑣𝑒 𝑏𝑖𝑎𝑠 = 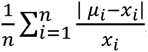, where *n* is the number of pseudo-observations, *xᵢ* the pseudo-observed parameter value, and *μᵢ* the posterior mode or mean for that replicate.

To assess the importance of LDD in shaping the gradient of Neanderthal ancestry in FA, we performed an additional set of 100,000 simulations, using the same simulation framework and parameter priors, but without using LDD. The fit of these simulations to the observed data was assessed in the same way described earlier for the analysis of the latitude of Neanderthal habitat.

Lastly, we performed an additional set of 100,000 simulations, using the same simulation framework and parameter priors as for the parameter estimation, simulating a scenario where the Neolithic transition was purely cultural. This means that no FA expansion was simulated and the FA samples were drawn directly from the HG layer. We used an ABC analysis to compare this scenario to the scenario that included the Neolithic expansion, to investigate if the observed difference in NE ancestry between the pre- and post-Neolithic European populations could be the result of drift alone or if the arrival of FA populations resulted in the dilution of NE ancestry.

## Acknowledgments

We thank André Langaney for carefully reading an initial version of the manuscript and providing valuable feedback.

## Funding

This work was supported by the Swiss National Science Foundation grants 31003A_182577 (to M.C.).

## Author contributions

Conceptualization: M.C., J.R., and A.T. Project administration: M.C. Data curation: A.T. and J.R., Methodology: A.T., M.C., J.R., and C.S.Q. Formal analysis: A.T. and J.R. Investigation: A.T. and J.R. Software: A.T. and M.C. Visualization: A.T. Writing—Original draft: A.T. Writing— Review and Editing: A.T., J.R., C.S.Q., and M.C. Resources: M.C. Validation: A.T. Supervision: M.C. Funding acquisition: M.C.

## Competing interests

The authors declare that they have no competing interests.

## Data and materials availability

All data needed to evaluate the conclusions in the paper are present in the paper and/or the Supplementary Materials. All genomes analyzed in this study were already published, and references are provided in data S1. The used version of SPLATCHE3 is provided in the online free repository Zenodo.org (https://zenodo.org/records/10495323). Example of input setting files can be shared upon request to the authors.

## Notes

### Competing Interest Statement

The authors have declared no competing interest.

## References

1. C. Stringer, The origin and evolution of Homo sapiens. Philosophical Transactions of the Royal Society B: Biological Sciences 371, 20150237 (2016).

2. A. Bergström, C. Stringer, M. Hajdinjak, E. M. L. Scerri, P. Skoglund, Origins of modern human ancestry. Nature 590, 229–237 (2021).

3. K. Prüfer, C. de Filippo, S. Grote, F. Mafessoni, P. Korlević, M. Hajdinjak, B. Vernot, L. Skov, P. Hsieh, S. Peyrégne, D. Reher, C. Hopfe, S. Nagel, T. Maricic, Q. Fu, C. Theunert, R. Rogers, P. Skoglund, M. Chintalapati, M. Dannemann, B. J. Nelson, F. M. Key, P. Rudan, Ž. Kućan, I. Gušić, L. V. Golovanova, V. B. Doronichev, N. Patterson, D. Reich, E. E. Eichler, M. Slatkin, M. H. Schierup, A. M. Andrés, J. Kelso, M. Meyer, S. Pääbo, A high-coverage Neandertal genome from Vindija Cave in Croatia. Science 358, 655–658 (2017).

4. J. Ochando, J. S. Carrión, R. Blasco, S. Fernández, G. Amorós, M. Munuera, P. Sañudo, J. Fernández Peris, Silvicolous Neanderthals in the far West: the mid-Pleistocene palaeoecological sequence of Bolomor Cave (Valencia, Spain). Quaternary Science Reviews 217, 247–267 (2019).

5. K. Prüfer, F. Racimo, N. Patterson, F. Jay, S. Sankararaman, S. Sawyer, A. Heinze, G. Renaud, P. H. Sudmant, C. de Filippo, H. Li, S. Mallick, M. Dannemann, Q. Fu, M. Kircher, M. Kuhlwilm, M. Lachmann, M. Meyer, M. Ongyerth, M. Siebauer, C. Theunert, A. Tandon, P. Moorjani, J. Pickrell, J. C. Mullikin, S. H. Vohr, R. E. Green, I. Hellmann, P. L. F. Johnson, H. Blanche, H. Cann, J. O. Kitzman, J. Shendure, E. E. Eichler, E. S. Lein, T. E. Bakken, L. V. Golovanova, V. B. Doronichev, M. V. Shunkov, A. P. Derevianko, B. Viola, M. Slatkin, D. Reich, J. Kelso, S. Pääbo, The complete genome sequence of a Neanderthal from the Altai Mountains. Nature 505, 43–49 (2014).

6. T. Higham, K. Douka, R. Wood, C. B. Ramsey, F. Brock, L. Basell, M. Camps, A. Arrizabalaga, J. Baena, C. Barroso-Ruíz, C. Bergman, C. Boitard, P. Boscato, M. Caparrós, N. J. Conard, C. Draily, A. Froment, B. Galván, P. Gambassini, A. Garcia-Moreno, S. Grimaldi, P. Haesaerts, B. Holt, M.-J. Iriarte-Chiapusso, A. Jelinek, J. F. Jordá Pardo, J.-M. Maíllo-Fernández, A. Marom, J. Maroto, M. Menéndez, L. Metz, E. Morin, A. Moroni, F. Negrino, E. Panagopoulou, M. Peresani, S. Pirson, M. de la Rasilla, J. Riel-Salvatore, A. Ronchitelli, D. Santamaria, P. Semal, L. Slimak, J. Soler, N. Soler, A. Villaluenga, R. Pinhasi, R. Jacobi, The timing and spatiotemporal patterning of Neanderthal disappearance. Nature 512, 306–309 (2014).

7. I. Djakovic, A. Key, M. Soressi, Optimal linear estimation models predict 1400–2900 years of overlap between Homo sapiens and Neandertals prior to their disappearance from France and northern Spain. Sci Rep 12, 15000 (2022).

8. R. E. Green, J. Krause, A. W. Briggs, T. Maricic, U. Stenzel, M. Kircher, N. Patterson, H. Li, W. Zhai, M. H.-Y. Fritz, N. F. Hansen, E. Y. Durand, A.-S. Malaspinas, J. D. Jensen, T. Marques-Bonet, C. Alkan, K. Prüfer, M. Meyer, H. A. Burbano, J. M. Good, R. Schultz, A. Aximu-Petri, A. Butthof, B. Höber, B. Höffner, M. Siegemund, A. Weihmann, C. Nusbaum, E. S. Lander, C. Russ, N. Novod, J. Affourtit, M. Egholm, C. Verna, P. Rudan, D. Brajkovic, Ž. Kucan, I. Gušic, V. B. Doronichev, L. V. Golovanova, C. Lalueza-Fox, M. de la Rasilla, J. Fortea, A. Rosas, R. W. Schmitz, P. L. F. Johnson, E. E. Eichler, D. Falush, E. Birney, J. C. Mullikin, M. Slatkin, R. Nielsen, J. Kelso, M. Lachmann, D. Reich, S. Pääbo, A Draft Sequence of the Neandertal Genome. Science 328, 710–722 (2010).

9. P. F. Reilly, A. Tjahjadi, S. L. Miller, J. M. Akey, S. Tucci, The contribution of Neanderthal introgression to modern human traits. Current Biology 32, R970–R983 (2022).

10. K. Harris, R. Nielsen, The Genetic Cost of Neanderthal Introgression. Genetics 203, 881–891 (2016).

11. I. Juric, S. Aeschbacher, G. Coop, The Strength of Selection against Neanderthal Introgression. PLOS Genetics 12, e1006340 (2016).

12. C. N. Simonti, B. Vernot, L. Bastarache, E. Bottinger, D. S. Carrell, R. L. Chisholm, D. R. Crosslin, S. J. Hebbring, G. P. Jarvik, I. J. Kullo, R. Li, J. Pathak, M. D. Ritchie, D. M. Roden, S. S. Verma, G. Tromp, J. D. Prato, W. S. Bush, J. M. Akey, J. C. Denny, J. A. Capra, The phenotypic legacy of admixture between modern humans and Neandertals. Science 351, 737–741 (2016).

13. H. Zeberg, The major genetic risk factor for severe COVID-19 is associated with protection against HIV. Proceedings of the National Academy of Sciences 119, e2116435119 (2022).

14. H. Zeberg, S. Pääbo, The major genetic risk factor for severe COVID-19 is inherited from Neanderthals. Nature 587, 610–612 (2020).

15. C. S. Quilodrán, J. Rio, A. Tsoupas, M. Currat, Past human expansions shaped the spatial pattern of Neanderthal ancestry. Science Advances 9, eadg9817 (2023).

16. C. S. Quilodrán, A. Tsoupas, M. Currat, The Spatial Signature of Introgression After a Biological Invasion With Hybridization. Frontiers in Ecology and Evolution 8 (2020).

17. L. N. Di Santo, C. S. Quilodrán, P. Cerrito, M. Currat, Tracing the Neanderthal–Modern Human hybrid zone using paleogenomic data. bioRxiv [Preprint] (2026). 10.64898/2026.01.06.697899.

18. M. Currat, M. Ruedi, R. J. Petit, L. Excoffier, The hidden side of invasions: Massive introgression by local genes. Evolution 62, 1908–1920 (2008).

19. M. E. Allentoft, M. Sikora, K.-G. Sjögren, S. Rasmussen, M. Rasmussen, J. Stenderup, P. B. Damgaard, H. Schroeder, T. Ahlström, L. Vinner, A.-S. Malaspinas, A. Margaryan, T. Higham, D. Chivall, N. Lynnerup, L. Harvig, J. Baron, P. D. Casa, P. Dąbrowski, P. R. Duffy, A. V. Ebel, A. Epimakhov, K. Frei, M. Furmanek, T. Gralak, A. Gromov, S. Gronkiewicz, G. Grupe, T. Hajdu, R. Jarysz, V. Khartanovich, A. Khokhlov, V. Kiss, J. Kolář, A. Kriiska, I. Lasak, C. Longhi, G. McGlynn, A. Merkevicius, I. Merkyte, M. Metspalu, R. Mkrtchyan, V. Moiseyev, L. Paja, G. Pálfi, D. Pokutta, Ł. Pospieszny, T. D. Price, L. Saag, M. Sablin, N. Shishlina, V. Smrčka, V. I. Soenov, V. Szeverényi, G. Tóth, S. V. Trifanova, L. Varul, M. Vicze, L. Yepiskoposyan, V. Zhitenev, L. Orlando, T. Sicheritz-Pontén, S. Brunak, R. Nielsen, K. Kristiansen, E. Willerslev, Population genomics of Bronze Age Eurasia. Nature 522, 167–172 (2015).

20. W. Haak, I. Lazaridis, N. Patterson, N. Rohland, S. Mallick, B. Llamas, G. Brandt, S. Nordenfelt, E. Harney, K. Stewardson, Q. Fu, A. Mittnik, E. Bánffy, C. Economou, M. Francken, S. Friederich, R. G. Pena, F. Hallgren, V. Khartanovich, A. Khokhlov, M. Kunst, P. Kuznetsov, H. Meller, O. Mochalov, V. Moiseyev, N. Nicklisch, S. L. Pichler, R. Risch, M. A. Rojo Guerra, C. Roth, A. Szécsényi-Nagy, J. Wahl, M. Meyer, J. Krause, D. Brown, D. Anthony, A. Cooper, K. W. Alt, D. Reich, Massive migration from the steppe was a source for Indo-European languages in Europe. Nature 522, 207–211 (2015).

21. Z. Hofmanová, S. Kreutzer, G. Hellenthal, C. Sell, Y. Diekmann, D. Díez-del-Molino, L. van Dorp, S. López, A. Kousathanas, V. Link, K. Kirsanow, L. M. Cassidy, R. Martiniano, M. Strobel, A. Scheu, K. Kotsakis, P. Halstead, S. Triantaphyllou, N. Kyparissi-Apostolika, D. Urem-Kotsou, C. Ziota, F. Adaktylou, S. Gopalan, D. M. Bobo, L. Winkelbach, J. Blöcher, M. Unterländer, C. Leuenberger, Ç. Çilingiroğlu, B. Horejs, F. Gerritsen, S. J. Shennan, D. G. Bradley, M. Currat, K. R. Veeramah, D. Wegmann, M. G. Thomas, C. Papageorgopoulou, J. Burger, Early farmers from across Europe directly descended from Neolithic Aegeans. PNAS 113, 6886–6891 (2016).

22. S. Shennan, The First Farmers of Europe: An Evolutionary Perspective (Cambridge University Press, Cambridge, 2018; https://www.cambridge.org/core/books/first-farmers-of-europe/7F3B7991A3527EFB8A43FDF89D98A9D1)*Cambridge World Archaeology*.

23. B. Bramanti, M. G. Thomas, W. Haak, M. Unterlaender, P. Jores, K. Tambets, I. Antanaitis-Jacobs, M. N. Haidle, R. Jankauskas, C.-J. Kind, F. Lueth, T. Terberger, J. Hiller, S. Matsumura, P. Forster, J. Burger, Genetic discontinuity between local hunter-gatherers and central Europe’s first farmers. Science 326, 137–140 (2009).

24. P. Skoglund, H. Malmström, M. Raghavan, J. Storå, P. Hall, E. Willerslev, M. T. P. Gilbert, A. Götherström, M. Jakobsson, Origins and Genetic Legacy of Neolithic Farmers and Hunter-Gatherers in Europe. Science 336, 466–469 (2012).

25. A. Tsoupas, C. S. Reyna-Blanco, C. S. Quilodrán, J. Blöcher, M. Brami, D. Wegmann, J. Burger, M. Currat, Local increases in admixture with hunter-gatherers followed the initial expansion of Neolithic farmers across continental Europe. Science Advances 11, eadq9976 (2025).

26. I. Mathieson, I. Lazaridis, N. Rohland, S. Mallick, N. Patterson, S. A. Roodenberg, E. Harney, K. Stewardson, D. Fernandes, M. Novak, K. Sirak, C. Gamba, E. R. Jones, B. Llamas, S. Dryomov, J. Pickrell, J. L. Arsuaga, J. M. B. de Castro, E. Carbonell, F. Gerritsen, A. Khokhlov, P. Kuznetsov, M. Lozano, H. Meller, O. Mochalov, V. Moiseyev, M. A. R. Guerra, J. Roodenberg, J. M. Vergès, J. Krause, A. Cooper, K. W. Alt, D. Brown, D. Anthony, C. Lalueza-Fox, W. Haak, R. Pinhasi, D. Reich, Genome-wide patterns of selection in 230 ancient Eurasians. Nature 528, 499–503 (2015).

27. M. E. Allentoft, M. Sikora, A. Refoyo-Martínez, E. K. Irving-Pease, A. Fischer, W. Barrie, A. Ingason, J. Stenderup, K.-G. Sjögren, A. Pearson, B. Sousa da Mota, B. Schulz Paulsson, A. Halgren, R. Macleod, M. L. S. Jørkov, F. Demeter, L. Sørensen, P. O. Nielsen, R. A. Henriksen, T. Vimala, H. McColl, A. Margaryan, M. Ilardo, A. Vaughn, M. Fischer Mortensen, A. B. Nielsen, M. Ulfeldt Hede, N. N. Johannsen, P. Rasmussen, L. Vinner, G. Renaud, A. Stern, T. Z. T. Jensen, G. Scorrano, H. Schroeder, P. Lysdahl, A. D. Ramsøe, A. Skorobogatov, A. J. Schork, A. Rosengren, A. Ruter, A. Outram, A. A. Timoshenko, A. Buzhilova, A. Coppa, A. Zubova, A. M. Silva, A. J. Hansen, A. Gromov, A. Logvin, A. B. Gotfredsen, B. Henning Nielsen, B. González-Rabanal, C. Lalueza-Fox, C. J. McKenzie, C. Gaunitz, C. Blasco, C. Liesau, C. Martinez-Labarga, D. V. Pozdnyakov, D. Cuenca-Solana, D. O. Lordkipanidze, D. En’shin, D. C. Salazar-García, T. D. Price, D. Borić, E. Kostyleva, E. V. Veselovskaya, E. R. Usmanova, E. Cappellini, E. Brinch Petersen, E. Kannegaard, F. Radina, F. Eylem Yediay, H. Duday, I. Gutiérrez-Zugasti, I. Merts, I. Potekhina, I. Shevnina, I. Altinkaya, J. Guilaine, J. Hansen, J. E. Aura Tortosa, J. Zilhão, J. Vega, K. Buck Pedersen, K. Tunia, L. Zhao, L. N. Mylnikova, L. Larsson, L. Metz, L. Yepiskoposyan, L. Pedersen, L. Sarti, L. Orlando, L. Slimak, L. Klassen, M. Blank, M. González-Morales, M. Silvestrini, M. Vretemark, M. S. Nesterova, M. Rykun, M. F. Rolfo, M. Szmyt, M. Przybyła, M. Calattini, M. Sablin, M. Dobisíková, M. Meldgaard, M. Johansen, N. Berezina, N. Card, N. A. Saveliev, O. Poshekhonova, O. Rickards, O. V. Lozovskaya, O. Gábor, O. C. Uldum, P. Aurino, P. Kosintsev, P. Courtaud, P. Ríos, P. Mortensen, P. Lotz, P. Persson, P. Bangsgaard, P. de Barros Damgaard, P. Vang Petersen, P. P. Martinez, P. Włodarczak, R. V. Smolyaninov, R. Maring, R. Menduiña, R. Badalyan, R. Iversen, R. Turin, S. Vasilyev, S. Wåhlin, S. Borutskaya, S. Skochina, S. A. Sørensen, S. H. Andersen, T. Jørgensen, Y. B. Serikov, V. I. Molodin, V. Smrcka, V. Merts, V. Appadurai, V. Moiseyev, Y. Magnusson, K. H. Kjær, N. Lynnerup, D. J. Lawson, P. H. Sudmant, S. Rasmussen, T. S. Korneliussen, R. Durbin, R. Nielsen, O. Delaneau, T. Werge, F. Racimo, K. Kristiansen, E. Willerslev, Population genomics of post-glacial western Eurasia. Nature 625, 301–311 (2024).

28. M. Currat, M. Arenas, C. S. Quilodràn, L. Excoffier, N. Ray, SPLATCHE3: simulation of serial genetic data under spatially explicit evolutionary scenarios including long-distance dispersal. Bioinformatics 35, 4480–4483 (2019).

29. M. A. Beaumont, W. Zhang, D. J. Balding, Approximate Bayesian Computation in Population Genetics. Genetics 162, 2025–2035 (2002).

30. S. Mallick, A. Micco, M. Mah, H. Ringbauer, I. Lazaridis, I. Olalde, N. Patterson, D. Reich, The Allen Ancient DNA Resource (AADR): A curated compendium of ancient human genomes. bioRxiv [Preprint] (2023). 10.1101/2023.04.06.535797.

31. M. Currat, L. Excoffier, Modern Humans Did Not Admix with Neanderthals during Their Range Expansion into Europe. PLOS Biology 2, e421 (2004).

32. T. K. Nielsen, B. M. Benito, J.-C. Svenning, B. Sandel, L. McKerracher, F. Riede, P. C. Kjærgaard, Investigating Neanderthal dispersal above 55°N in Europe during the Last Interglacial Complex. Quaternary International 431, 88–103 (2017).

33. M. Weiss, M. Hein, B. Urban, M. C. Stahlschmidt, S. Heinrich, Y. H. Hilbert, R. C. Power, H. v. Suchodoletz, T. Terberger, U. Böhner, F. Klimscha, S. Veil, K. Breest, J. Schmidt, D. Colarossi, M. Tucci, M. Frechen, D. C. Tanner, T. Lauer, Neanderthals in changing environments from MIS 5 to early MIS 4 in northern Central Europe – Integrating archaeological, (chrono)stratigraphic and paleoenvironmental evidence at the site of Lichtenberg. Quaternary Science Reviews 284, 107519 (2022).

34. C. Finlayson, F. Giles Pacheco, J. Rodríguez-Vidal, D. A. Fa, J. María Gutierrez López, A. Santiago Pérez, G. Finlayson, E. Allue, J. Baena Preysler, I. Cáceres, J. S. Carrión, Y. Fernández Jalvo, C. P. Gleed-Owen, F. J. Jimenez Espejo, P. López, J. Antonio López Sáez, J. Antonio Riquelme Cantal, A. Sánchez Marco, F. Giles Guzman, K. Brown, N. Fuentes, C. A. Valarino, A. Villalpando, C. B. Stringer, F. Martinez Ruiz, T. Sakamoto, Late survival of Neanderthals at the southernmost extreme of Europe. Nature 443, 850–853 (2006).

35. W. E. Banks, F. d’Errico, A. T. Peterson, M. Kageyama, A. Sima, M.-F. Sánchez-Goñi, Neanderthal Extinction by Competitive Exclusion. PLOS ONE 3, e3972 (2008).

36. C. Stringer, P. Andrews, The Complete World of Human Evolution (Thames & Hudson, London, Second Edition., 2012; https://www.thamesandhudsonusa.com/books/the-complete-world-of-human-evolution-softcover-second-edition).

37. M. Melchionna, M. Di Febbraro, F. Carotenuto, L. Rook, A. Mondanaro, S. Castiglione, C. Serio, V. A. Vero, G. Tesone, M. Piccolo, J. A. F. Diniz-Filho, P. Raia, Fragmentation of Neanderthals’ pre-extinction distribution by climate change. Palaeogeography, Palaeoclimatology, Palaeoecology 496, 146–154 (2018).

38. L. Slimak, T. Vimala, A. Seguin-Orlando, L. Metz, C. Zanolli, R. Joannes-Boyau, M. Frouin, L. J. Arnold, M. Demuro, T. Devièse, D. Comeskey, M. Buckley, H. Camus, X. Muth, J. E. Lewis, H. Bocherens, P. Yvorra, C. Tenailleau, B. Duployer, H. Coqueugniot, O. Dutour, T. Higham, M. Sikora, Long genetic and social isolation in Neanderthals before their extinction. Cell Genomics 4 (2024).

39. O. François, M. Currat, N. Ray, E. Han, L. Excoffier, J. Novembre, Principal Component Analysis under Population Genetic Models of Range Expansion and Admixture. Molecular Biology and Evolution 27, 1257–1268 (2010).

40. J.-J. Hublin, N. Sirakov, V. Aldeias, S. Bailey, E. Bard, V. Delvigne, E. Endarova, Y. Fagault, H. Fewlass, M. Hajdinjak, B. Kromer, I. Krumov, J. Marreiros, N. L. Martisius, L. Paskulin, V. Sinet-Mathiot, M. Meyer, S. Pääbo, V. Popov, Z. Rezek, S. Sirakova, M. M. Skinner, G. M. Smith, R. Spasov, S. Talamo, T. Tuna, L. Wacker, F. Welker, A. Wilcke, N. Zahariev, S. P. McPherron, T. Tsanova, Initial Upper Palaeolithic Homo sapiens from Bacho Kiro Cave, Bulgaria. Nature 581, 299–302 (2020).

41. P. Mellars, The earliest modern humans in Europe. Nature 479, 483–485 (2011).

42. M. Currat, L. Excoffier, The effect of the Neolithic expansion on European molecular diversity. Proceedings of the Royal Society of London B: Biological Sciences 272, 679–688 (2005).

43. M. Petr, S. Paabo, J. Kelso, B. Vernot, Limits of long-term selection against Neandertal introgression. P Natl Acad Sci USA 116, 1639–1644 (2019).

44. D. N. Vyas, C. J. Mulligan, Analyses of Neanderthal introgression suggest that Levantine and southern Arabian populations have a shared population history. American Journal of Physical Anthropology 169, 227–239 (2019).

45. B. Vernot, J. M. Akey, Complex History of Admixture between Modern Humans and Neandertals. The American Journal of Human Genetics 96, 448–453 (2015).

46. L. Chen, A. B. Wolf, W. Fu, L. Li, J. M. Akey, Identifying and Interpreting Apparent Neanderthal Ancestry in African Individuals. Cell 180, 677–687.e16 (2020).

47. J. D. Wall, M. A. Yang, F. Jay, S. K. Kim, E. Y. Durand, L. S. Stevison, C. Gignoux, A. Woerner, M. F. Hammer, M. Slatkin, Higher Levels of Neanderthal Ancestry in East Asians than in Europeans. Genetics 194, 199–209 (2013).

48. K. E. Witt, F. Villanea, E. Loughran, X. Zhang, E. Huerta-Sanchez, Apportioning archaic variants among modern populations. Philosophical Transactions of the Royal Society B: Biological Sciences 377, 20200411 (2022).

49. M. Currat, L. Excoffier, Strong reproductive isolation between humans and Neanderthals inferred from observed patterns of introgression. Proceedings of the National Academy of Sciences of the United States of America 108, 15129–15134 (2011).

50. L. Excoffier, C. S. Quilodrán, M. Currat, “Models of hybridization during range expansions and their application to recent human evolution” in Cultural Developments in the Eurasian Paleolithic and the Origin of Anatomically Modern Humans, A. P. Derevianko, M. Shunkov, Eds. (Department of the Institute of Archaeology and Ethnography SB RAS, Novosibirsk, Russia, 2014), pp. 122–137.

51. R. E. Green, A.-S. Malaspinas, J. Krause, A. W. Briggs, P. L. F. Johnson, C. Uhler, M. Meyer, J. M. Good, T. Maricic, U. Stenzel, K. Prüfer, M. Siebauer, H. A. Burbano, M. Ronan, J. M. Rothberg, M. Egholm, P. Rudan, D. Brajković, Ž. Kućan, I. Gušić, M. Wikström, L. Laakkonen, J. Kelso, M. Slatkin, S. Pääbo, A Complete Neandertal Mitochondrial Genome Sequence Determined by High-Throughput Sequencing. Cell 134, 416–426 (2008).

52. A. W. Briggs, J. M. Good, R. E. Green, J. Krause, T. Maricic, U. Stenzel, C. Lalueza-Fox, P. Rudan, D. Brajković, Ž. Kućan, I. Gušić, R. Schmitz, V. B. Doronichev, L. V. Golovanova, M. de la Rasilla, J. Fortea, A. Rosas, S. Pääbo, Targeted Retrieval and Analysis of Five Neandertal mtDNA Genomes. Science 325, 318–321 (2009).

53. F. Mafessoni, S. Grote, C. De Filippo, V. Slon, K. A. Kolobova, B. Viola, S. V. Markin, M. Chintalapati, S. Peyrégne, L. Skov, P. Skoglund, A. I. Krivoshapkin, A. P. Derevianko, M. Meyer, J. Kelso, B. Peter, K. Prüfer, S. Pääbo, A high-coverage Neandertal genome from Chagyrskaya Cave. Proc. Natl. Acad. Sci. U.S.A. 117, 15132–15136 (2020).

54. L. Li, T. J. Comi, R. F. Bierman, J. M. Akey, Recurrent gene flow between Neanderthals and modern humans over the past 200,000 years. Science 385, eadi1768 (2024).

55. H. Li, R. Durbin, Inference of human population history from individual whole-genome sequences. Nature 475, 493–6 (2011).

56. B. Vanmontfort, Forager–farmer connections in an ‘unoccupied’ land: First contact on the western edge of LBK territory. Journal of Anthropological Archaeology 27, 149–160 (2008).

57. A. Kousathanas, C. Leuenberger, V. Link, C. Sell, J. Burger, D. Wegmann, Inferring Heterozygosity from Ancient and Low Coverage Genomes. Genetics 205, 317–332 (2017).

58. C. E. G. Amorim, T. Hofer, N. Ray, M. Foll, A. Ruiz-Linares, L. Excoffier, Long-distance dispersal suppresses introgression of local alleles during range expansions. Heredity 118, 135–142 (2017).

59. M. Tallavaara, M. Luoto, N. Korhonen, H. Järvinen, H. Seppä, Human population dynamics in Europe over the Last Glacial Maximum. PNAS 112, 8232–8237 (2015).

60. A. Timmermann, Quantifying the potential causes of Neanderthal extinction: Abrupt climate change versus competition and interbreeding. Quaternary Science Reviews 238, 106331 (2020).

61. A. Degioanni, C. Bonenfant, S. Cabut, S. Condemi, Living on the edge: Was demographic weakness the cause of Neanderthal demise? PLOS ONE 14, e0216742 (2019).

62. C. N. Roberts, J. Woodbridge, A. Palmisano, A. Bevan, R. Fyfe, S. Shennan, Mediterranean landscape change during the Holocene: Synthesis, comparison and regional trends in population, land cover and climate. The Holocene 29, 923–937 (2019).

63. A. Timpson, S. Colledge, E. Crema, K. Edinborough, T. Kerig, K. Manning, M. G. Thomas, S. Shennan, Reconstructing regional population fluctuations in the European Neolithic using radiocarbon dates: a new case-study using an improved method. Journal of Archaeological Science 52, 549–557 (2014).

64. N. M. Silva, S. Kreutzer, A. Souleles, S. Triantaphyllou, K. Kotsakis, D. Urem-Kotsou, P. Halstead, N. Efstratiou, S. Kotsos, G. Karamitrou-Mentessidi, F. Adaktylou, A. Chondroyianni-Metoki, M. Pappa, C. Ziota, A. Sampson, A. Papathanasiou, K. Vitelli, T. Cullen, N. Kyparissi-Apostolika, A. Z. Lanz, J. Peters, J. Rio, D. Wegmann, J. Burger, M. Currat, C. Papageorgopoulou, Ancient mitochondrial diversity reveals population homogeneity in Neolithic Greece and identifies population dynamics along the Danubian expansion axis. Sci Rep 12, 13474 (2022).

65. A. P. Sümer, H. Rougier, V. Villalba-Mouco, Y. Huang, L. N. M. Iasi, E. Essel, A. Bossoms Mesa, A. Furtwaengler, S. Peyrégne, C. de Filippo, A. B. Rohrlach, F. Pierini, F. Mafessoni, H. Fewlass, E. I. Zavala, D. Mylopotamitaki, R. A. Bianco, A. Schmidt, J. Zorn, B. Nickel, A. Patova, C. Posth, G. M. Smith, K. Ruebens, V. Sinet-Mathiot, A. Stoessel, H. Dietl, J. Orschiedt, J. Kelso, H. Zeberg, K. I. Bos, F. Welker, M. Weiss, S. P. McPherron, T. Schüler, J.-J. Hublin, P. Velemínský, J. Brůžek, B. M. Peter, M. Meyer, H. Meller, H. Ringbauer, M. Hajdinjak, K. Prüfer, J. Krause, Earliest modern human genomes constrain timing of Neanderthal admixture. Nature 638, 711–717 (2025).

66. N. Marchi, L. Winkelbach, I. Schulz, M. Brami, Z. Hofmanová, J. Blöcher, C. S. Reyna-Blanco, Y. Diekmann, A. Thiéry, A. Kapopoulou, V. Link, V. Piuz, S. Kreutzer, S. M. Figarska, E. Ganiatsou, A. Pukaj, T. J. Struck, R. N. Gutenkunst, N. Karul, F. Gerritsen, J. Pechtl, J. Peters, A. Zeeb-Lanz, E. Lenneis, M. Teschler-Nicola, S. Triantaphyllou, S. Stefanović, C. Papageorgopoulou, D. Wegmann, J. Burger, L. Excoffier, The genomic origins of the world’s first farmers. Cell 185, 1842–1859.e18 (2022).

67. V. Villalba-Mouco, M. S. van de Loosdrecht, A. B. Rohrlach, H. Fewlass, S. Talamo, H. Yu, F. Aron, C. Lalueza-Fox, L. Cabello, P. Cantalejo Duarte, J. Ramos-Muñoz, C. Posth, J. Krause, G.-C. Weniger, W. Haak, A 23,000-year-old southern Iberian individual links human groups that lived in Western Europe before and after the Last Glacial Maximum. Nat Ecol Evol 7, 597–609 (2023).

68. J. Rio, C. S. Quilodrán, M. Currat, Spatially explicit paleogenomic simulations support cohabitation with limited admixture between Bronze Age Central European populations. Commun Biol 4, 1–12 (2021).

69. S. Urbanek, proj4: A simple interface to the PROJ.4 cartographic projections library, version 1.0-12 (2022); https://cran.r-project.org/web/packages/proj4/index.html.

70. R. J. Wang, S. I. Al-Saffar, J. Rogers, M. W. Hahn, Human generation times across the past 250,000 years. Science Advances 9, eabm7047 (2023).

71. C. Stringer, L. Crété, Mapping Interactions of H. neanderthalensis and Homo sapiens from the Fossil and Genetic Records. PaleoAnthropology 2022 (2022).

72. P. Mellars, J. C. French, Tenfold Population Increase in Western Europe at the Neandertal–to–Modern Human Transition. Science 333, 623–627 (2011).

73. M. Kimura, “Stepping stone” model of population. Annual Report of the National Institute of Genetics Japan 3, 62–63 (1953).

74. N. Nakatsuka, É. Harney, S. Mallick, M. Mah, N. Patterson, D. Reich, ContamLD: estimation of ancient nuclear DNA contamination using breakdown of linkage disequilibrium. Genome Biol 21, 199 (2020).

75. D. Reich, K. Thangaraj, N. Patterson, A. L. Price, L. Singh, Reconstructing Indian population history. Nature 461, 489–494 (2009).

76. R. Maier, P. Flegontov, O. Flegontova, U. Işıldak, P. Changmai, D. Reich, On the limits of fitting complex models of population history to f-statistics. eLife 12, e85492 (2023).

77. J. Pinheiro, B. Douglas, D. Saikat, S. Deepayan, EISPACK autohrs, H. Siem, V. W. Bert, R. Johannes, R Core Team, nlme: Linear and Nonlinear Mixed Effects Models, version 3.1-164 (2023); https://cran.r-project.org/web/packages/nlme/index.html.

78. W. Roebroeks, N. J. Conard, T. van Kolfschoten, R. W. Dennell, R. C. Dunnell, C. Gamble, P. Graves, K. Jacobs, M. Otte, D. Roe, J. Svoboda, A. Tuffreau, B. A. Voytek, F. Wenban-Smith, J. J. Wymer, Dense Forests, Cold Steppes, and the Palaeolithic Settlement of Northern Europe [and Comments and Replies]. Current Anthropology 33, 551–586 (1992).

79. T. K. Nielsen, F. Riede, On Research History and Neanderthal Occupation at its Northern Margins. European Journal of Archaeology 21, 506–527 (2018).

80. D. Wegmann, C. Leuenberger, S. Neuenschwander, L. Excoffier, ABCtoolbox: a versatile toolkit for approximate Bayesian computations. BMC Bioinformatics 11, 116 (2010).

81. P. Pudlo, J.-M. Marin, A. Estoup, J.-M. Cornuet, M. Gautier, C. P. Robert, Reliable ABC model choice via random forests. Bioinformatics 32, 859–866 (2016).

82. J. Novembre, M. Stephens, Interpreting principal component analyses of spatial population genetic variation. Nat Genet 40, 646–649 (2008).

83. C. Leuenberger, D. Wegmann, Bayesian Computation and Model Selection Without Likelihoods. Genetics 184, 243–252 (2010).

