## Supplementary Materials for "Tracing Neanderthal ancestry patterns through successive population expansions in Europe"

Alexandros Tsoupas *et al.*

**This PDF file includes:**

Figs. S1 to S4
Tables S1 to S5

**Other Supplementary Materials for this manuscript include the following:**

Legends for data S1 to S2

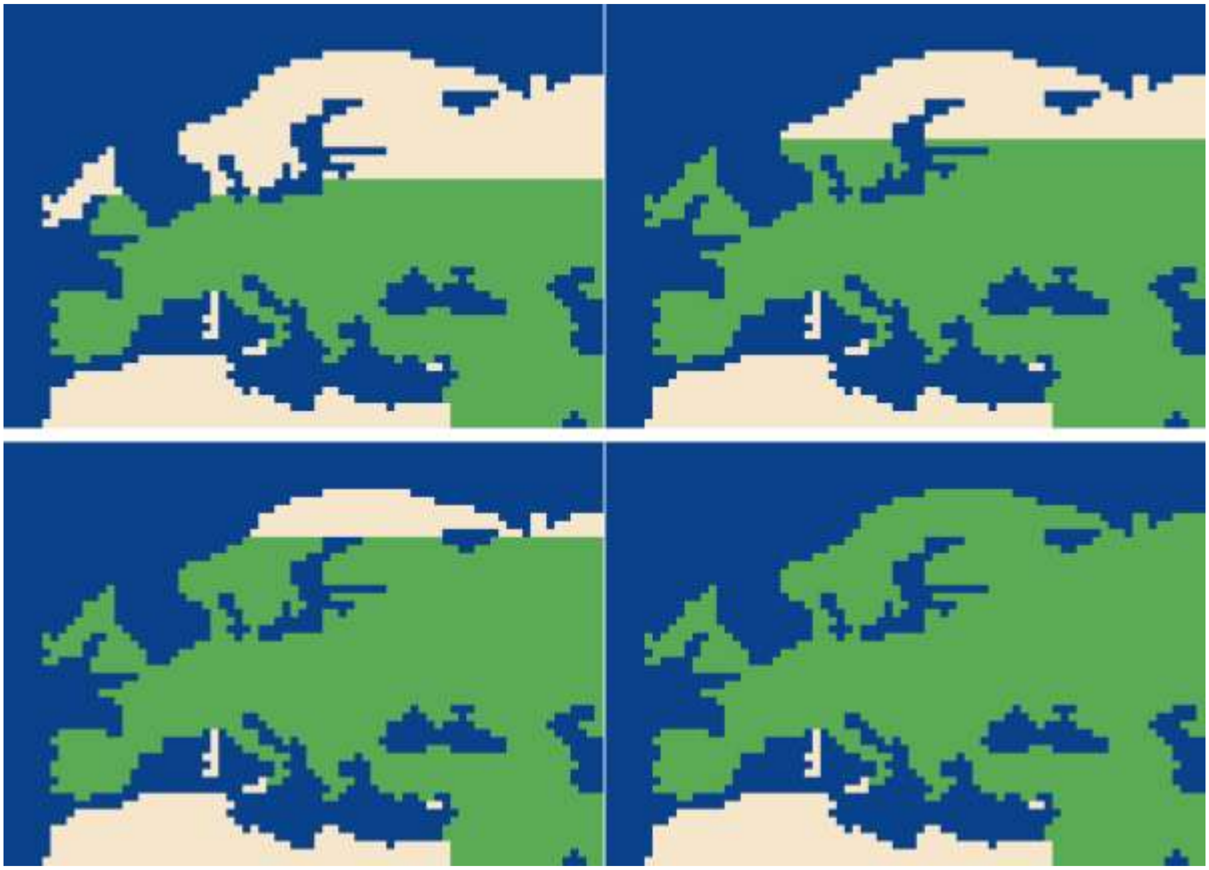

**Supplementary Figure S1. The four tested northern limits of Neanderthal habitat.** Green areas indicate regions habitable by Neanderthals, white those considered uninhabitable. Scenario names are indicated in parentheses in each panel.

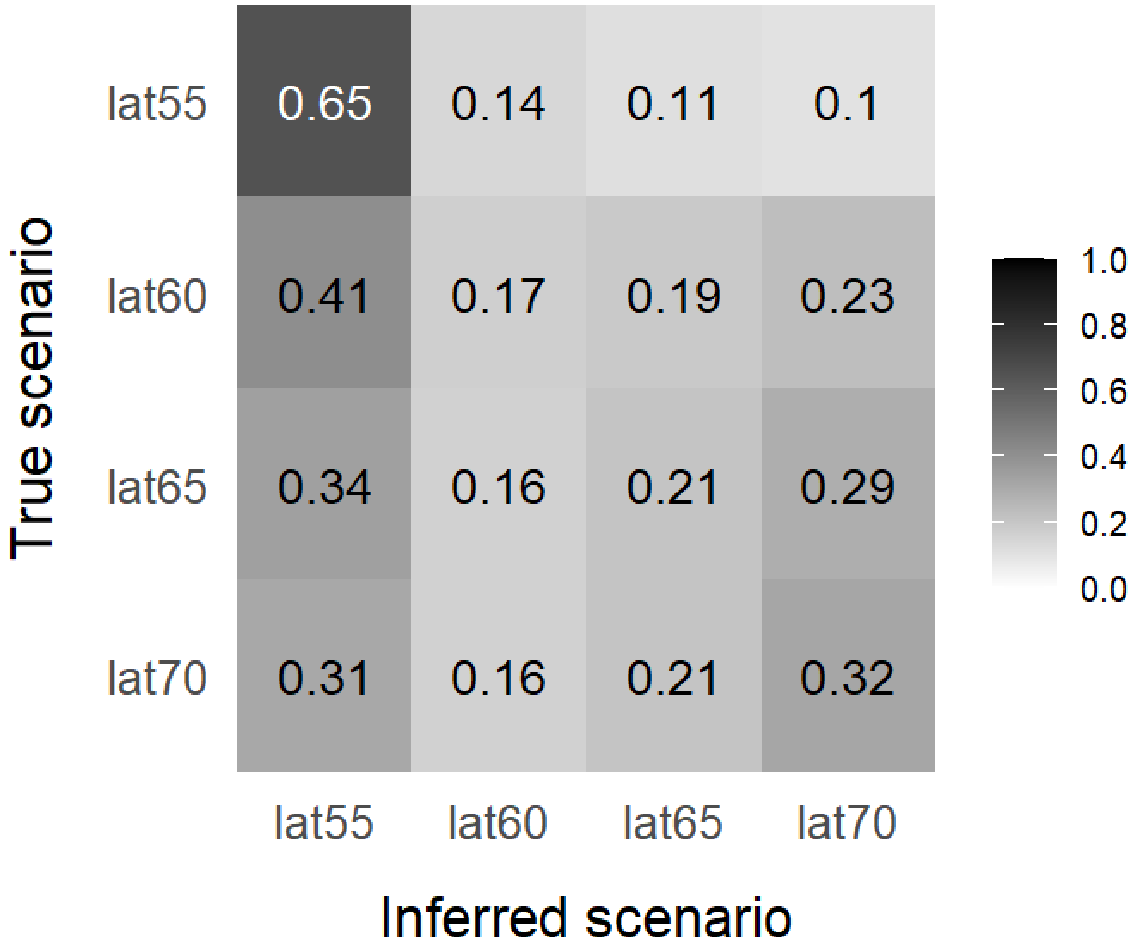


**Supplementary Figure S2. Confusion matrix of the four investigated scenarios of Neanderthal habitat northern latitudinal limits.** Calculation performed with the abcrf R package (ABC-random forest approach; *81*), using 2,000 trees and 400,000 simulations per scenario. Each row represents simulations from a given scenario (the 'true' scenario), while the columns indicate the proportion of those simulations that ABC attributed to each of the studied scenarios (the 'inferred' scenarios).

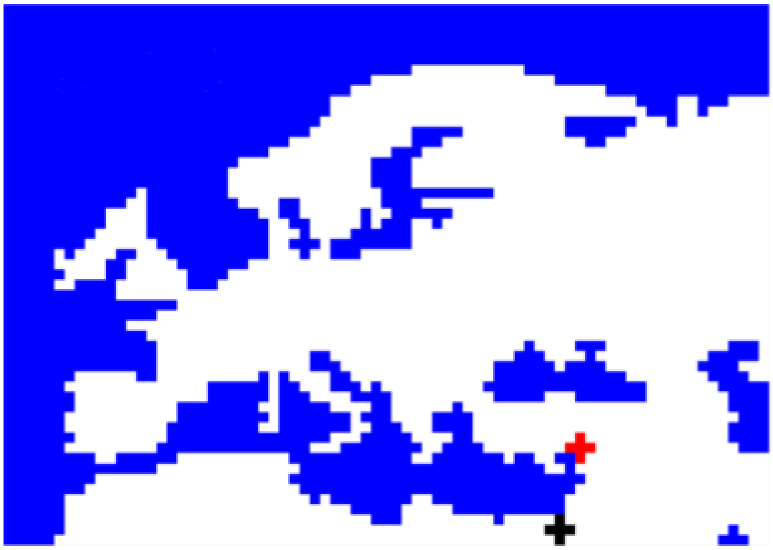

**Supplementary Figure S3**. **The locations chosen as starting points for the successive AMH expansions in the simulations for parameter estimation** (Black = Hunter-gatherer (HG) source, Red = Farmer (FA) source).

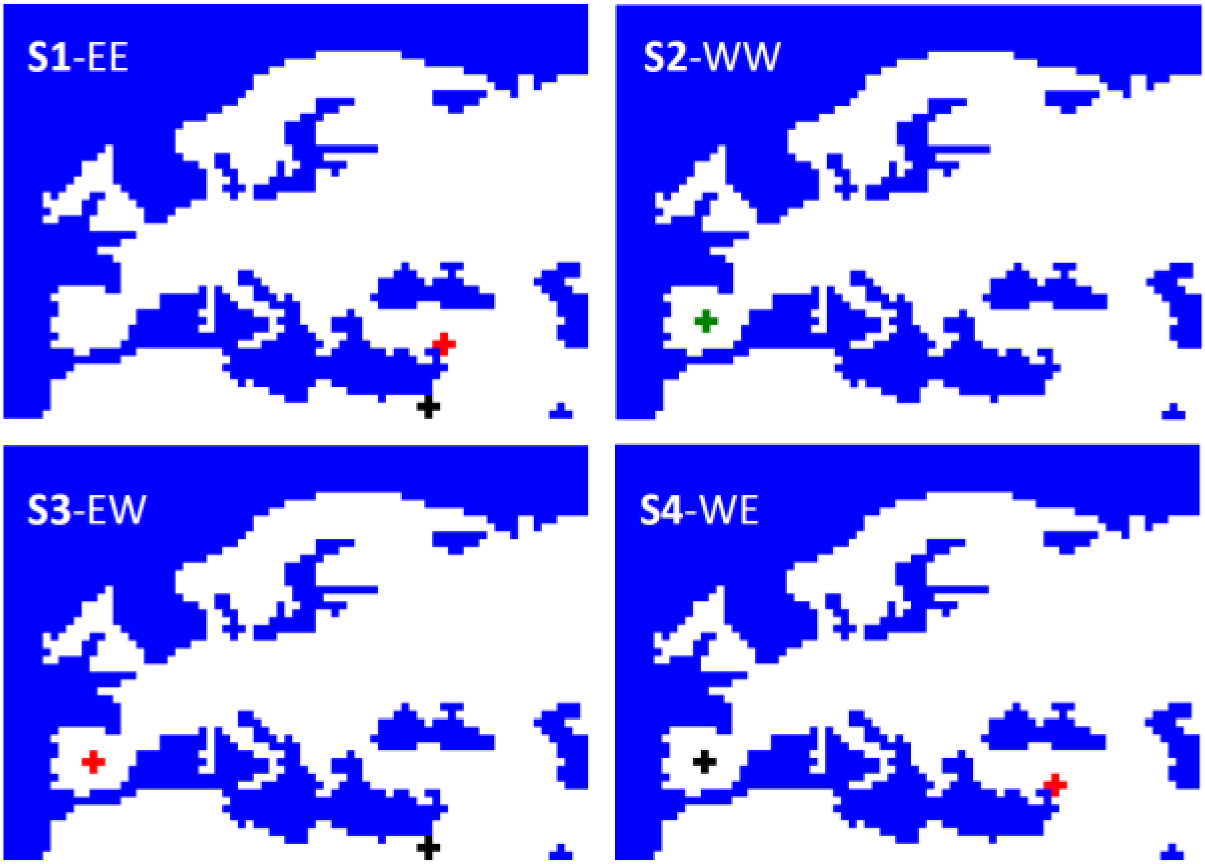


**Supplementary Figure S4. The four expansion scenarios tested, defined by the starting locations of HG and FA.** Black crosses mark HG origins, red crosses FA origins, and green crosses a shared origin. Scenario names are indicated in each panel.

**Supplementary Table S1. Marginal density *p*-values for the simulated statistics of the four scenarios investigating the Neanderthal habitat latitudinal limits.** Estimated on 400,000 simulations per scenario with three tolerance levels, retaining 2,000, 4,000 or 12,000 simulations. Computed with ABCtoolbox2 (*80*).

| Tolerance level | Marginal posterior density p-value of scenario lat55 | Marginal posterior density p-value of scenario lat60 | Marginal posterior density p-value of scenario lat65 | Marginal posterior density p-value of scenario lat70 |
|---|---|---|---|---|
| 0.005 | 0.614 | 0.364 | 0.280 | 0.277 |
| 0.01 | 0.523 | 0.273 | 0.234 | 0.242 |
| 0.03 | 0.568 | 0.324 | 0.231 | 0.188 |

**Supplementary Table S2. Model choice between the four investigated latitudinal limits of Neanderthal habitat.** Performed with abcrf R package (ABC-random forest approach; *81*), using 2,000 trees and 400,000 simulations per scenario. The votes refer to the number of trees that selected each scenario as the most probable one.

| Scenario lat55 | Scenario lat60 | Scenario lat65 | Scenario lat70 | Chosen scenario | Posterior probability of chosen scenario |
|---|---|---|---|---|---|
| 1068 | 447 | 254 | 231 | lat55 | 0.56 |

**Supplementary Table S3. Proportion of simulations of the four investigated scenarios of northern Neanderthal presence, which produced a gradient of Neanderthal ancestry in HG increasing with distance from the expansion source, a gradient of NE ancestry decreasing with time or both these gradients.**

| Scenario | Proportion of simulations producing spatially increasing NE ancestry gradient in HG | Proportion of simulations producing temporally decreasing NE ancestry gradient in HG | Proportion of simulations producing both spatial and temporal gradients in HG |
|---|---|---|---|
| lat55 | 0.80 | 0.66 | 0.63 |
| lat60 | 0.82 | 0.55 | 0.54 |
| lat65 | 0.82 | 0.50 | 0.49 |
| lat70 | 0.82 | 0.47 | 0.46 |

**Supplementary Table S4. Mean difference in Neanderthal (NE) ancestry between hunter-gatherers (HG) and farmers (FA) across simulated scenarios with different expansion origins.** For each simulation, the difference was calculated as mean Neanderthal ancestry in HG minus that in FA, and results were then averaged across all simulations of each scenario.

| Scenario | Mean NE ancestry difference between HG and FA | Welch's *t*-test *p*-value |
|---|---|---|
| S1-EE | 0.0065 | $<10^{-07}$ |
| S2-WW | 0.0258 | $<10^{-07}$ |
| S3-EW | -0.0214 | $<10^{-07}$ |
| S4-WE | -0.0087 | $<10^{-07}$ |

**Supplementary Table S5. Characteristics of the posterior distributions of the estimated parameters.** The estimation was performed on 200,000 simulations of the 3-layered approach, with three tolerance levels, retaining 1,000, 2,000 or 6,000 simulations. Computed with ABCtoolbox2 (*80*). NE stands for Neanderthals, HG for Paleolithic hunter-gatherers and FA for Neolithic farmers.

| Parameter | Tolerance level | Posterior mode | Posterior mode relative bias | Posterior mean | Posterior mean relative bias | Posterior lower HDI95 | Posterior upper HDI95 |
|---|---|---|---|---|---|---|---|
| Hybridization rate between NE and HG ($\gamma_H$) | 0.005 | 0.0058 | 0.087 | 0.0058 | 0.087 | 0.0039 | 0.0079 |
| | **0.01** | **0.006** | **0.088** | **0.0059** | **0.088** | **0.0038** | **0.008** |
| | 0.03 | 0.0062 | 0.105 | 0.0061 | 0.105 | 0.0039 | 0.0083 |
| Assimilation rate of HG into FA populations ($\gamma_A$) | 0.005 | 0.0444 | 1.515 | 0.0906 | 1.733 | 0.001 | 0.1789 |
| | **0.01** | **0.0485** | **1.931** | **0.0919** | **2.214** | **0.0019** | **0.1808** |
| | 0.03 | 0.0424 | 2.448 | 0.0863 | 3.212 | 0 | 0.1762 |
| Ratio of HG to NE effective population size ($Kratio_{HN}$) | 0.005 | 6.444 | 0.442 | 6.094 | 0.385 | 2.387 | 9.798 |
| | **0.01** | **7.657** | **0.484** | **6.088** | **0.401** | **2.315** | **9.717** |
| | 0.03 | 7.495 | 0.787 | 6.000 | 0.614 | 2.283 | 9.670 |
| Ration of FA to HG effective population size ($Kratio_{FH}$) | 0.005 | 3.849 | 0.230 | 4.924 | 0.221 | 3.101 | 6.779 |
| | **0.01** | **4.253** | **0.221** | **4.943** | **0.220** | **3.135** | **6.818** |
| | 0.03 | 4.535 | 0.216 | 5.002 | 0.216 | 3.183 | 6.859 |
| Proportion of Long Distance Dispersals ($P_{LDD}$) | 0.005 | 0.0015 | 1.89 | 0.0032 | 4.489 | 0 | 0.008 |
| | **0.01** | **0.0015** | **1.813** | **0.0032** | **4.320** | **0** | **0.0078** |
| | 0.03 | 0.0015 | 1.823 | 0.0033 | 4.404 | 0 | 0.008 |

**Data S1. (separate file)**

**Information of the genomic data used in the present study.** The information provided are: the name assigned to the individual (AADR_VersionID); the study which produced the genome (Publication); if the individual was classified as being a hunter-gatherer (HG) or a farmer (FA) (Population_Group); the latitude of the area where the individual was found (Latitude); the longitude of the area where the individual was found (Longitude); the age of the individual in years before present (Mean_Age_YBP); the generation at which the individual is sampled dusting the simulations (Simulated_generation); the population sample to which the individual was assigned for estimating the mean sample Neanderthal ancestry and for performing the simulations (Sample); the estimated Neanderthal ancestry of the individual.

**Data S2. (separate file)**

**Information of the population samples into which the individuals were grouped.** The information provided are: the name of the sample (Sample_name); if the sample was sampled from the layer of hunter-gatherers (HG) or farmers (FA) (Population_Group); the latitude of the deme where the sample was simulated (Latitude); the longitude of the deme where the sample was simulated (Longitude); the distance of the sample deme form the respective expansion source in *Km* (Distance_from_expansion_source_in_km); the mean age of the individuals in the sample, in years before present (Mean_sample_age_YBP); the generation at which the sample is sampled during the simulations (Simulated_generation); the population sample to which the individual was assigned for estimating the mean sample Neanderthal ancestry and for performing the simulations (Sample); the for FA samples, the sub-division of the Neolithic to which the sample was assigned for linear mixed model analysis (Simulated_period); the mean estimated Neanderthal ancestry of the individuals in the sample (Mean_Neanderthal_ancestry).
